# Genomic forecasts of maladptation in *Lycaeides* butterflies

**DOI:** 10.64898/2026.05.16.725655

**Authors:** Kenen B. Goodwin, Samridhi Chaturvedi, Lauren K. Lucas, Zachariah Gompert

## Abstract

Genomic forecasting approaches based on genotype-environment associations (GEAs) are increasingly used to estimate genomic offsets (GOs), which predict population maladaptation and extinction risk under current or future climatic conditions. Despite their widespread use, only a subset of studies have evaluated how accurately GOs predict (mal)adaptation, limiting their interpretation and application in policy and management. Here, we used GEA analyses to estimate GOs for past, present, and future climates in *Lycaeides* butterflies, focusing on the causes of variation in GOs among populations and their relationships with demographic parameters inferred from population genomic data. Using multivariate linear regression and genotyping-by-sequencing data from 42 *Lycaeides* populations (922 butterflies), we found that mean annual temperature, cumulative annual precipitation, and hybridization history together explained 47.6% of variation in genome-wide allele frequencies. Genomic offsets differed substantially among populations and across past, present, and future climates, with evidence for increasing maladaptation under more distant future climate scenarios. We found no relationship between GOs for present climates and contemporary effective population size. In contrast, genetic diversity, which reflects long-term effective population size, and local rates of gene flow together explained 27.3% of variation in contemporary GOs. Populations with higher genetic diversity and more gene flow exhibited lower GOs, consistent with the hypothesis that genetic diversity enhances adaptive capacity and that gene flow may introduce adaptive alleles. Overall, our results support the utility of GO predictions, particularly when validated with independent measures of adaptation, while cautioning against simplistic interpretations of GO as a direct measure of maladaptation in conservation and management contexts.

## Introduction

Organisms are often well adapted to the environments they occupy, but rapid environmental change can shift fitness optima fast enough to generate maladaptation (Schluter, 2000; Schlaepfer *et al*., 2002; Hereford, 2009; Brady *et al*., 2019; Capblancq *et al*., 2020). Such mal-adaptation can lead to demographic decline and even extinction if populations fail to adapt rapidly or if organisms are unable to disperse to more suitable habitats (Orr & Unckless, 2008; Bell & Gonzalez, 2011; Travis *et al*., 2013; R^ego *et al*., 2019; Marrec & Bitbol, 2020; Morgan *et al*., 2020; Anstett *et al*., 2026; Wu *et al*., 2026). Indeed, evidence of maladaptation, population decline, and extinction resulting from rapid anthropogenic climate change and habitat alteration is widespread (McLaughlin *et al*., 2002; Parmesan, 2006; Maclean & Wilson, 2011; Wiens, 2016; Halsch *et al*., 2021; Edwards *et al*., 2025). Predicting which populations or species are most likely to become maladapted, and thus face an elevated risk of decline or extinction, is therefore a major goal in biology (Bernatchez *et al*., 2024).

Much past work on predicting persistence in novel or altered environments has relied on species distribution modeling (e.g., Bakkenes *et al*., 2002; Thomas *et al*., 2004; Jezkova & Wiens, 2016). These approaches generally estimate a species’ climatic niche from occurrence data and use the inferred niche to forecast the species’ range under predicted future climatic conditions (Elith & Leathwick, 2009). Species distribution models have been successful in some cases (Hijmans & Graham, 2006; Tingley *et al*., 2009), but their failure to account for genetic and phenotypic variation among populations can lead to over- or underestimation of extinction risk (Fitzpatrick & Keller, 2015; Exposito-Alonso *et al*., 2018; Rellstab *et al*., 2021). Such information can be obtained from common garden experiments, which predict (mal)adaptation of genotypes and phenotypes in novel environments by measuring the performance or fitness of organisms under controlled conditions that mimic future environments of interest (Pelini *et al*., 2012; Exposito-Alonso *et al*., 2019; Morente-Ĺopez *et al*., 2022). Thus, common garden experiments complement species distribution models and are particularly powerful for detecting adaptation and predicting performance, although they can be difficult or costly to implement for many species.

More recently, a diverse family of genomic forecasting approaches has been developed to predict the adaptive and demographic consequences of climate change (Capblancq *et al*., 2020; Rellstab *et al*., 2021; Bernatchez *et al*., 2024). These methods account for adaptive genetic differences among populations without the resource-intensive requirements of common garden experiments, making them applicable to a wider range of organisms. Numerous approaches and metrics for genomic prediction of maladaptation have been introduced, including genomic offset (Capblancq *et al*., 2020), gradient forest genetic offset (Fitzpatrick & Keller, 2015), risk of nonadaptedness (RONA) (Rellstab *et al*., 2016), and genetic gap (Gain *et al*., 2023). Most of these metrics have well-defined mathematical relationships with one another (Gain *et al*., 2023), and we hereafter use genomic offset (GO) in a broad sense to refer to this collection of approaches.

Genomic offset methods first estimate associations between multivariate environmental variables and multilocus genotypes or allele frequencies (Capblancq *et al*., 2020; Capblancq & Forester, 2021). The estimated genotype–environment associations (GEAs) are then used to calculate (i) the deviation of a population’s (or individual’s) allele frequencies (or genotypes) from the predicted optimum for the current environment (current GO), or (ii) the deviation between the predicted optima for the current and a future environment (future GO) (Borrell *et al*., 2020; Capblancq *et al*., 2020; Gain *et al*., 2023). Under the assumption that GEAs reflect local adaptation and that predicted allele frequencies correspond to optimal allele frequencies, these current or future GOs quantify the magnitude of multilocus allele frequency change required for adaptation to an environment or maintenance of adaptedness in a future environment, and by extension the degree of maladaptation to the current or future environment (Fitzpatrick & Keller, 2015; Borrell *et al*., 2020; Rellstab *et al*., 2021). In other words, populations with larger GOs are considered at greater risk of decline or extinction under current or future climates. Recent theoretical and empirical work, however, has shown that the success of genomic forecasts is context-dependent, being influenced by the biology of the system, the goals of the prediction, the evaluation method, and the novelty of the target environment (Lind *et al*., 2024; Lotterhos, 2024; Fitzpatrick *et al*., 2026).

Genomic offset approaches have now been applied in many systems, particularly trees (Borrell *et al*., 2020; Ingvarsson & Bernhardsson, 2020; Fitzpatrick *et al*., 2021; Hung *et al*., 2023; Yuan *et al*., 2023), but also in animals such as birds and mammals (Bay *et al*., 2018b; Chen *et al*., 2022; Teng *et al*., 2025). In contrast, GO methods have not been widely adopted for insects, despite their crucial role in most ecosystems and their status as one of the animal groups most affected by climate change (Halsch *et al*., 2021; Harvey *et al*., 2023; Forister *et al*., 2024; Edwards *et al*., 2025). Moreover, relatively few studies have tested or validated genomic predictions of (mal)adaptation (Hoffmann *et al*., 2021; Rellstab *et al*., 2021). Some validations do exist, most notably in predicting the growth and survival of trees planted in common garden experiments (e.g., Borrell *et al*., 2020; Fitzpatrick *et al*., 2021; Lind *et al*., 2024; Verrico *et al*., 2026). Results from these studies have generally been favorable, suggesting that GO metrics are better predictors of performance in common gardens than alternative measures, such as the geographic or climatic distance between the source population and the common garden. However, these successes may not directly translate into valid predictions for demographic trends in nature, which depend on survival, dispersal, and reproduction (Lotterhos, 2024).

Some studies have begun connecting GO metrics to demographic outcomes. For example, genomic predictions of future climate vulnerability in the yellow warbler (*Setophaga petechia*) were moderately predictive of recent population declines based on breeding bird surveys (Bay *et al*., 2018b). This validation links past demographic trends to future predictions rather than past predictions, but is nonetheless promising (Fitzpatrick *et al*., 2018; Bay *et al*., 2018a). Thus, while there are reasons for cautious optimism, GOs should not be interpreted automatically as direct measures of genomic vulnerability to maladaptation. Much remains unknown about the predictive power of GO metrics and what these metrics reveal across diverse systems (Lotterhos, 2024; Fitzpatrick *et al*., 2026). Our study contributes to filling this knowledge gap.

Here, we generate and analyze genomic predictions of climatic (mal)adaptation in *Lycaeides* butterflies. Our goal is twofold. First, we estimate GOs in terms of current (mal)adaptation to past, present, and future climatic conditions based on precipitation and temperature, in order to assess evidence of adaptation across space and time. Second, we test the relationship between GOs and genetic estimates of demographic parameters that may correlate with adaptive capacity, specifically contemporary effective population size (related to current population size), genetic diversity (indicative of long-term effective population size), and gene flow (i.e., the degree of connectivity among populations). In doing so, we treat these demographic correlates not as direct measures of fitness, but as an independent axis of validation that helps clarify what GOs may and may not capture in natural populations. We focus on these parameters (i.e., population size and population connectivity) because they can be readily inferred for most systems using the same data employed to estimate GOs. We find that GOs vary among populations and time periods, with stronger evidence of maladaptation to future climates, and that GOs appear to track genetic diversity and connectivity more closely than contemporary effective population size. More generally, our work suggests that GOs may contain information relevant to conservation, but also cautions against simplistic interpretations of GOs as direct estimates of population decline or local extinction (i.e., genomic vulnerability). Our results further highlight the need for multifaceted validation approaches before applying GOs in conservation and management contexts.

## Methods

### Study system

*Lycaeides* butterflies exist in North America as a complex of four or more closely related nominal species and numerous recognized subspecies (Scott, 1992; Gompert *et al*., 2014). This group includes the federally endangered Karner blue butterfly (*Lycaeides samuelis*) (Gompert *et al*., 2006; Forister *et al*., 2011) and the Lotis blue butterfly (*Lycaeides anna lotis*), which was last seen in the wild in the 1983 and is now presumed extinct (Arnold, 1980, 1993). Here, we focus on two species, *L. idas* and *L. melissa*, that occur throughout western North America where their ranges partially overlap (Nabokov, 1943; Gompert *et al*., 2010, 2014). These species exhibit subtle differences in male genital morphology (Nabokov, 1944; Lucas *et al*., 2008; Gompert *et al*., 2010), wing pattern elements (Lucas *et al*., 2018), host plant use (Nice *et al*., 2002; Gompert *et al*., 2013) and voltinism (Gompert *et al*., 2013). Despite these differences, past and ongoing hybridization between these *L. idas* and *L. melissa* is common, resulting in a broad cline in genetic ancestry in the central Rocky Mountains (Gompert *et al*., 2010, 2012, 2014; Chaturvedi *et al*., 2020; Zhang *et al*., 2023). We thus treat this pair of species and the genetically intervening populations as an entity (i.e., a species complex) for the current study (we incorporate genetic ancestry in our analyses).

### DNA sequence data, alignment, and variant calling

We obtained previously published DNA sequence data from 42 *Lycaeides* butterfly populations, with a mean of 22 individuals per population (range = 13–46, 922 total, see Table 1 for details) (Gompert *et al*., 2014; Chaturvedi *et al*., 2018). These data comprise 896,277,495 100 base pair (bp) Illumina sequences from genotyping-by-sequencing (GBS) libraries (following Parchman *et al*., 2012; Gompert *et al*., 2014). We aligned the DNA sequences to a chromosome-level reference genome for *L. melissa* (Zhang *et al*., 2023). We did this using the bwa mem algorithm (version 0.7.19-r1273) (Li & Durbin, 2009). We then compressed, filtered and indexed the sequence alignments with samtools (version 1.16) (Li *et al*., 2009). Next, we identified genetic variants (SNPs) using the consensus variant caller from bcftools (version 1.16) (Li *et al*., 2009). For this, we excluded reads with mapping qualities *<*20 and bases with quality scores *<*30, ignored indels, and only designated SNPs when the posterior probability of a nucleotide being invariant was less than 0.01 with the default prior variant probability of 0.001. We then filtered the initial set of putative SNPs to retain only those SNPs with a minimum average read depth of 2*→* per individual (1844*→* across all samples), at least 10 reads supporting the non-reference allele, a quality score of 30, missing data (no reads) for fewer than 20% of individuals, no other SNP within 3 bps, and coverage that did not exceed the mean across SNPs by more than three standard deviations. This left us with 142,307 SNPs for population genomic analyses.

**Table 1:** Summary of populations analyzed in this study, including population identifiers, population names, longitude and latitude (WGS84), the data source (Data) for DNA sequences (A = Gompert *et al*., 2014, B = Chaturvedi *et al*., 2018), the mean admixture proportion from entropy for each population (*q̅*), and the cumulative annual precipitation and mean annual temperature data used in the genotype-environment-association analyses. An admixture proportion of 1 represents *Lycaeides melissa*, whereas an admixture proportion of 0 represents *Lycaeides idas*.

| ID | Population name | Long.° | Lat.° | Data | $\bar{q}$ | Precip. (mm) | Temp. (°C) |
| --- | --- | --- | --- | --- | --- | --- | --- |
| ABC | Abel Creek, NV | -117.619 | 41.425 | B | 0.926 | 353.063 | 9.865 |
| ABM | Albion Meadows, UT | -111.623 | 40.587 | A | 0.668 | 1240.994 | 3.535 |
| BIC | Big Ice Cave, WY | -108.401 | 45.162 | A | 0.222 | 565.337 | 4.035 |
| BHP | Bishop, CA | -118.391 | 37.358 | A | 1.000 | 122.125 | 13.730 |
| BTB | Blacktail Butte, WY | -110.682 | 43.638 | A | 0.158 | 578.500 | 3.395 |
| BSD | Brandon, SD | -96.582 | 43.605 | B | 0.751 | 725.558 | 7.800 |
| BCR | Bull Creek, WY | -110.553 | 43.301 | A | 0.180 | 649.470 | 2.560 |
| BNP | Bunsen Peak, WY | -110.721 | 44.934 | A | 0.047 | 507.969 | 2.620 |
| CDY | Cody, WY | -108.978 | 44.512 | A | 0.688 | 269.695 | 7.410 |
| CKV | Cokeville, WY | -110.937 | 42.005 | A | 0.940 | 307.463 | 4.375 |
| VCP | Crystal Peak Park, NV | -119.995 | 39.514 | B | 0.985 | 405.562 | 10.210 |
| DBQ | De Beque, CO | -108.209 | 39.321 | A | 0.968 | 273.540 | 11.175 |
| DCR | Deeth-Charleston, NV | -115.376 | 41.302 | A | 0.937 | 359.705 | 6.025 |
| GVL | Gardnerville, NV | -119.779 | 38.812 | A | 0.997 | 436.457 | 9.895 |
| GNP | Garnet Peak, MT | -111.225 | 45.432 | A | 0.031 | 599.396 | 4.400 |
| GLA | Goose Lake, CA | -120.293 | 41.986 | A | 0.972 | 491.325 | 7.935 |
| HNV | Hayden Valley, WY | -110.495 | 44.682 | A | 0.059 | 652.559 | 0.895 |
| KHL | King's Hill, MT | -110.699 | 46.841 | A | 0.019 | 964.115 | 1.470 |
| LCA | Lamoille Canyon, NV | -115.471 | 40.679 | A | 0.955 | 516.721 | 6.020 |
| LAN | Lander, WY | -108.355 | 42.653 | A | 0.669 | 306.705 | 6.875 |
| MTU | Montague, CA | -122.377 | 41.772 | A | 1.000 | 660.708 | 10.455 |
| MON | Montrose, CO | -107.816 | 38.375 | A | 0.967 | 289.811 | 8.895 |
| OCY | Ophir City, NV | -117.274 | 38.944 | A | 0.988 | 401.584 | 6.700 |
| PSP | Periodic Spring, WY | -110.849 | 42.747 | A | 0.211 | 909.314 | 3.075 |
| PIN | Pinnacles, WY | -109.976 | 43.740 | A | 0.206 | 905.213 | 0.100 |
| REW | Red Earth Way, NV | -118.836 | 38.981 | A | 0.999 | 119.776 | 11.640 |
| RNV | Rendezvous Mt., WY | -110.885 | 43.596 | A | 0.173 | 1449.364 | 0.085 |
| RDL | Riddle Lake, WY | -110.546 | 44.362 | A | 0.185 | 773.248 | 0.930 |
| SHC | Shovel Creek, CA | -122.164 | 41.881 | A | 0.001 | 495.272 | 9.095 |
| SVY | Sierra Valley, CA | -120.369 | 39.643 | B | 1.000 | 583.204 | 8.455 |
| SLA | Silver Lake, NV | -119.927 | 39.650 | A | 0.999 | 282.941 | 10.075 |
| SYC | Siyeh Creek, MT | -113.668 | 48.703 | A | 0.001 | 2347.201 | 2.130 |
| SDC | Soldier Creek, MT | -114.612 | 47.206 | A | 0.035 | 505.514 | 5.970 |
| SCC | Star Creek Canyon, NV | -118.123 | 40.547 | A | 0.984 | 365.613 | 9.400 |
| SUV | Surprise Valley, CA | -120.099 | 41.282 | A | 0.952 | 348.620 | 8.965 |
| TIB | Tibb's Butte, WY | -109.452 | 44.950 | A | 0.093 | 988.839 | -1.160 |
| TBY | Tomboy Road, CO | -107.774 | 37.942 | A | 0.024 | 882.384 | 1.830 |
| TPT | Trout Pond Trailhead, CA | -116.579 | 32.977 | B | 1.000 | 719.986 | 13.200 |
| UAL | Upper Alkali Lake, CA | -120.173 | 41.793 | B | 0.963 | 445.486 | 8.800 |
| VIC | Victor, ID | -111.111 | 43.659 | A | 0.854 | 529.837 | 4.600 |
| WAL | Washoe Lake, NV | -119.779 | 39.233 | A | 1.000 | 337.276 | 9.890 |
| YWP | Yellow Pine, WY | -105.401 | 41.252 | A | 0.729 | 546.348 | 4.615 |

### Estimating allele frequencies and admixture proportions

We estimated population allele frequencies, which were then used for the GEA analyses and to estimate GOs. We obtained maximum likelihood estimates of allele frequencies using estpEM (version 0.1) (Gompert *et al*., 2014), which implements the expectation-maximization (EM) algorithm described in Li (2011). This approach works directly with the genotype likelihoods from bcftools and thereby incorporates uncertainty in genotypes when estimating population allele frequencies. We set the convergence tolerance to 0.001 and allowed for a maximum of 50 EM iterations for this analysis.

We next estimated admixture proportions to describe the main axis of population structure in this data set (Gompert *et al*., 2014) and thus account for this as a latent covariate in the GEA analyses. Admixture proportions were inferred using entropy (version 1.2) (Gompert *et al*., 2014; Shastry *et al*., 2021) and assuming two source populations (i.e., *k* = 2), consistent with known patterns of structure and the major ancestry cline documented in the *L. idas*-*L. melissa* species complex (Gompert *et al*., 2010, 2014; Chaturvedi *et al*., 2020). The admixture proportion model in entropy is similar to the correlated allele frequencies admixture model in structure (Pritchard *et al*., 2000), but has an added feature of incorporating uncertainty in genotypes (as captured by genotype likelihoods) into estimates of admixture proportions. We estimated admixture proportions from a set of 5000 randomly selected common variants (SNPs with global minor allele frequency *>* 0.05) (see Figure S1). Bayesian estimates of admixture proportions were based on nine independent Markov chain Monte Carlo (MCMC) runs each comprising 2000 iterations and a 1000 iteration burnin with a thinning interval of 5. We calculated the Gelman-Rubin potential scale reduction factor to verify likely convergence of the MCMC algorithm to the posterior and adequate mixing (mean = 1.1, median = 1.1).

### Genotype-environment associations and genomic offset

To identify genetic loci associated with environmental conditions, we fit a multivariate-normal linear regression with population allele frequencies as the response variables and climate normals as predictors. Specifically, we hypothesized that *Lycaeides* butterflies may be sensitive to variations in temperature and precipitation, given the well-documented effects of these factors on butterfly abundance and phenology (Forister *et al*., 2023; Edwards *et al*., 2025; Reis *et al*., 2026). We also included admixture proportion as an additional predictor to control for differences in genetic composition associated with evolutionary history. To represent climate predictors, we extracted 30-year normals of mean annual temperature and cumulative annual precipitation from the 2020 PRISM climate normals (1991-2020 normals, 800-m resolution, version M4) (Daly *et al*., 1994). To assess potential interactions between temperature, precipitation, and admixture, we generated all three-and two-way interactions as additional covariates. All predictors were mean-centered and scaled by one standard deviation prior to the derivation of interaction terms.

We fit the multivariate-normal linear regression in R (version 4.2.0) (R Core Team, 2022) using the lm function, and we performed variable selection by comparing adjusted *R*^2^ values (Ezekiel, 1930) between competing models containing all combinations of predictors. We performed variable selection by adjusted *R*^2^ because computational errors in the multivariate-normal density function at high dimensionality precluded the use of information criteria. Further, adjusted *R*^2^ values are readily obtained for redundancy analysis (RDA) (Rao, 1964; Wollenberg, 1977) in the vegan package (version 2.7-2) (Oksanen *et al*., 2022). Because a multivariate-normal regression forms the base of an RDA, our modeling and variable selection approach are consistent with those of the widely-applied RDA for genotype-environment associations (e.g., Capblancq & Forester, 2021; Gain *et al*., 2023). Further, because principal component analysis (PCA) rotations of the response and fitted values within an RDA preserve both the total and explained variance, adjusted *R*^2^ values between a multivariate-normal linear regression and an RDA are identical (Legendre & Legendre, 2012). As such, we omitted the PCA rotations applied by RDA because these were not relevant to our analysis.

We took the multivariate-normal regression with the highest adjusted *R*^2^ value as the best-fitting model, and we performed all subsequent predictions and analyses using this best-fit model. To assess the contribution of each covariate to explaining variation in population allele frequencies, we calculated partial *R*^2^ values (Anderson-Sprecher, 1994) for each predictor in the best-fit model. A partial *R*^2^ value represents the percent reduction in unexplained variance when the predictor is included in the model (i.e., best-fit model), compared to a model without the predictor (i.e., best-fit model with predictor omitted). Under the assumption that model predictions represent optimal allele frequencies for a given climate and genetic background, we calculated GO as the Euclidean distance between observed population allele frequencies and their predicted values according to the best-fit model. Thus, GO is an estimate of how maladapted a population is to its environment. Because PCA rotations within an RDA preserve Euclidean distance (Legendre & Legendre, 2012), our approach to calculating GO is equivalent to that of conventional RDA applications (Gain *et al*., 2023).

### Genomic offset projections

For each population, we calculated GOs to past, present and future climatic conditions using the best-fit multivariate-normal regression model. In GO calculations, expected allele frequencies for current conditions were predicted using the same 30-year PRISM climate normals used to fit the multivariate-normal regression. Meanwhile, future climatic conditions were estimated from RegCM3 regionally-downscaled climate projections (Hostetler *et al*., 2011). The RegCM3 dataset contains downscaled projections of the A2 emissions scenario from several competing general circulation models (GCMs) with different forecast periods. Of the two included forecasts which extend to the end of the century (i.e., 2011-2099), we opted to use the forecast based on the GENMOM GCM (Alder *et al*., 2011) – which is both newer than the alternative ECHAM5 GCM (Roeckner *et al*., 2003) and has been applied extensively in climate research (Hostetler *et al*., 2011). From a conservation planning perspective, forecasts of the A2 emissions scenario are beneficial in providing an upper-intermediate expectation of climatic shifts (IPCC, 2007). As some populations of *Lycaeides* butterflies are federally-listed under the Endangered Species Act (USFWS, 1992), we believe that the A2 emissions scenario is appropriate for GO predictions of *Lycaeides* populations.

To account for differences in climate values between data sources (i.e., PRISM and RegCM3), we considered PRISM normals to be the “true” values for contemporary conditions, and we added decadal changes from within the RegCM3 dataset to contemporary PRISM normals to produce future climate projections for each population. We calculated decadal climate values within the RegCM3 dataset as the average of all monthly values within the preceding decade. For example, the decadal RegCM3 value for 2020 was the average of monthly values from 2011 to 2020. This calculation was repeated for each decade from 2020 to 2090, and the 2020 decadal value was subtracted from all later decades to estimate the projected change in temperature and precipitation since 2020. We completed our estimation of future climatic conditions by adding the 2020 PRISM climate normals to the RegCM3 decadal differences, producing a time series of temperature and precipitation projections spanning through 2090.

Past climatic conditions were estimated from PRISM climate estimates. Specifically, we extracted monthly climate estimates from 1981 to 2020 using the freely-available 4-km resolution dataset (version M3). Because the M3 version dataset only models precipitation back to 1981, we only consider climatic conditions after 1980 in our analyses. We generated past climate estimates using the same approach as future estimates. That is, monthly values were averaged across decadal intervals (e.g., 1981-1990), and their differences from the 2011-2020 interval were added to the 2020 PRISM climate normals. Hereafter, each decade is denoted by its final year (e.g., 1981-1990 as “1990”). For each population, the resulting time series of mean annual temperature and cumulative annual precipitation spans one century from 1990 to 2090 at decadal intervals. Using this time series, we generated GO predictions for each population during each decade using the best-fit model.

To examine temporal trends in genomic offset projections, we fit a linear mixed-effects model with year (fixed effect) and population (random effect) as predictors of log-transformed GO. The linear mixed-effects model was fit using the lmer function in the lme4 package (version 1.1.38) (Bates *et al*., 2015) to optimize the restricted maximum likelihood, and *p*-values for regression coefficients were generated through computation of Satterthwaite degrees of freedom (Satterthwaite, 1946) with the lmerTest package (version 3.2.0) (Kuznetsova *et al*., 2017). Under the hypothesis that *Lycaeides* genotypes represent local adaptation to recent climates, we predicted that GO would increase through time with increasing dissimilarity between projected and historic climatic conditions.

### Gene enrichment by environmental association

To assess the functional significance of genotype-environment associations, we performed gene enrichment tests for each predictor to determine whether the proportion of statistically-significant SNPs which were gene-associated was higher than expected from random sampling. For each predictor, SNPs were considered statistically-significant if their false-discovery-rate-adjusted *p*-values (Benjamini & Hochberg, 1995) were ≤ 0.05 in the best-fit multivariate-normal regression model. SNPs were considered gene-associated if functional annotations described the SNP as on or within 10,000 bases of a gene-coding sequence. We performed gene enrichment tests for each predictor in the best-fit model, in which one million iterations were used to generate each null distribution.

Within an iteration, we randomly drew the number of statistically-significant SNPs from the overall pool of SNPs in the study. For each enrichment test, we then calculated the *p*-value as the proportion of iterations in which the number of randomly-selected SNPs which were gene-associated matched or exceeded the number of statistically-significant SNPs which were gene-associated. Using the same methods, we also performed enrichment tests for gene ontology annotations. We used 10,000 iterations to generate each null distribution, and a false-discovery-rate adjustment was applied to all *p*-values within each predictor’s set of gene ontology enrichment tests. The gene ontology enrichment tests aimed to identify biological processes which were significantly associated with climatic conditions.

### Relationship between GOs and demographic factors

We next asked whether variation in GOs could be explained by genetic estimates of demographic factors, specifically contemporary effective population size, relative measures of genetic diversity (indicative of long-term effective population size), and effective gene flow (a measure of population connectivity). Contemporary (inbreeding) effective population sizes (herein denoted *N_e_*) were inferred based on patterns of linkage disequilibrium (LD) between unlinked loci (i.e., pairs of SNPs on different chromosomes) (Waples, 2024). We calculated LD using posterior estimates of genotypes from entropy (specifically, posterior means rounded to the nearest integer) for a set of 5000 common variants, excluding Z-linked loci. LD was measured as the squared genotypic correlation: 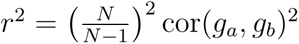, where *N* is the sample size for the population, and *g_a_* and *g_b_* are vectors of integer-valued genotypes for loci *a* and *b* (Waples & Do, 2008; Zaykin *et al*., 2008). This formulation is mathematically equivalent to the *r*^2^ standardization of Burrow’s Δ (i.e., a composite disequilibrium metric) (Weir, 1996; Zaykin, 2004; Gao *et al*., 2008; Waples, 2024).

For each population, we obtained an average LD measure by randomly sampling 10,000 pairs of unlinked SNPs from our dataset (this approach avoids the potential bias of including linked loci, see, e.g., Waples *et al*., 2016; Waples, 2024). This procedure was repeated 100 times to generate a distribution of estimates accounting for sub-sampling variability. LD estimates were corrected for the effect of sampling and then converted into estimates of *N_e_* following Waples (2006) (see Table S1). All analyses were conducted in R. Estimates of contemporary *N_e_* based on time-series data were available for a subset of seven populations (Gompert *et al*., 2021) and were highly correlated with the LD-based estimates of contemporary *N_e_* (Pearson correlation, *r* = 0.91, 95% CI = 0.52–0.99, *P* = 0.004), providing additional support that the LD-based metrics accurately capture local effective population sizes.

We used eems (version 0.0.0.9000) to infer effective migration rates and effective diversity levels for the 42 *Lycaeides* populations (Petkova *et al*., 2016). Analyses were based on posterior genotype estimates from entropy for 5000 randomly selected common SNPs. Rather than estimating absolute migration rates, this method identifies geographic regions with relatively low or high gene flow compared to a simple two-dimensional stepping-stone isolation-by-distance model. As a result, the approach can pinpoint areas experiencing limited or elevated dispersal or gene flow relative to null spatial expectations (Petkova *et al*., 2016). The effective diversity level parameters capture the average genetic dissimilarity between individuals sampled from the same deme and are expected to correlate with long-term effective population sizes, which inherently include the effects of gene flow on diversity. The model was fit using MCMC with three chains, each consisting of six million sampling iterations, three million burn-in iterations, and a thinning interval of 10,000. We fit the models twice, assuming either 400 or 500 demes evenly distributed on a triangular grid, and then averaged parameter estimates across the three chains and two assumed numbers of demes (see Figure S2).

We next considered whether GOs for present climate could be explained by our population genetic correlates of demographic parameters. Specifically, we fit a multiple linear regression between 2020 genomic offset and contemporary effective population size, effective migration, and effective diversity. We hypothesized that maladapted populations would exhibit lower *N_e_* than populations that were well-adapted to contemporary conditions. We further hypothesized that gene flow would contribute potentially-adaptive alleles, and we hypothesized that genetic diversity would provide adaptive capacity. Given these hypotheses, we expected a negative relationship between genomic offset and each of the three demographic parameters. We log-transformed the 2020 genomic offset response to improve the normality of residuals, and we used the median bootstrap *N_e_* estimate for each population. All covariates were mean-centered and scaled by one standard deviation. The Pearson correlation coefficients (*ρ*) for all pairwise combinations of predictors were low. *ρ* was highest between *N_e_* and effective diversity (0.199), and *ρ* was lowest between effective migration and effective diversity (0.022). *ρ* between *N_e_* and effective migration was also low at 0.063.

Variable selection was performed across all combinations of predictors using the expected log pointwise predictive density (elppd) estimated using leave-one-out cross-validation (LOO-CV). elppd via LOO-CV is an estimate of a model’s out-of-sample performance, as expressed via a predictive log-likelihood (Gelman *et al*., 2013). As Akaike’s information criterion (AIC) (Akaike, 1973) is an asymptotic approximation of cross-validation (Stone, 1977), elppd provides a more direct approach to model selection. To put elppd on the familiar deviance scale (as expressed by information criteria), we multiplied LOO-CV elppd values by negative two (Gelman *et al*., 2013). We considered the best-fitting model as the one with the lowest −2 × LOO-CV elppd, and we calculated partial *R*^2^ statistics for the predictors included in the best-fit model.

## Results

### Genotype-environment associations and genomic offset

By adjusted *R*^2^, the best-fit multivariate-normal linear regression model between allele frequencies and climatic conditions included all main effects of admixture, temperature, and precipitation. The best-fit model also included the three-way interaction between main effects, along with 2 two-way interactions involving admixture (i.e., admixture-temperature, admixture-precipitation). The best-fit model had an *R*^2^ value of 0.476, and an adjusted *R*^2^ value of 0.387. Many competing models had comparable adjusted *R*^2^ values (50% of models had adjusted *R*^2^ values greater than 0.3), and summaries for all competing models are presented in Table S2. Because our objective was to generate predictions of GO, we focus only on the best-fit model here. Partial *R*^2^ values were highest for admixture (0.229), followed by temperature (0.092) and their interaction (0.073). The lowest partial *R*^2^ values were associated with precipitation (0.030) and its interactions (Table S3). To quantify the overall explanatory contribution of climatic predictors, we generalized the concept of *R*^2^ to model comparison (Anderson-Sprecher, 1994). By treating the best-fit GEA as the full model and the admixture-only GEA as the reduced model, we computed a generalized *R*^2^ of 0.225, which suggests that – after controlling for the effect of admixture proportion – the climatic predictors and their interactions explained 22.5% of the remaining variation in population allele frequencies.

To examine whether climatic adaptations were concentrated in specific areas of the genome, we visualized FDR-adjusted *p*-values for each combination of predictor and SNP in the best-fit multivariate-normal linear regression model across chromosomal positions (Figure S3). For three predictors, chromosome Z contained the highest proportion of significantly-associated SNPs (i.e., FDR-adjusted *p*-values ≤ 0.05; admixture = 1.51%, precipitation = 0.42%, admixture-temperature-precipitation interaction = 0.35%). For the remaining three predictors, chromosome Z contained the second-highest proportion of significantly-associated SNPs behind chromosome 1 (temperature = 0.66% [chr. 1] & 0.51% [chr. Z], admixture-temperature interaction = 0.45% [chr. 1] & 0.38% [chr. Z], admixture-precipitation interaction = 0.17% [chr. 1] & 0.13% [chr. Z]). Thus, chromosomes Z and 1 had the highest concentrations of SNPs associated with climatic predictors, with chromosome Z containing either the highest or second-highest proportion of significant SNPs across all predictors in the best-fit model.

### Genomic offset projections

GO predictions from the best-fit multivariate-normal linear regression from 1990 through 2090 for each population are shown in Figures 1 and 2. Averaged across all populations, mean annual temperature was projected to increase by 2.964°C (σ = 0.385°C) between 1990 and 2090. On average, cumulative annual precipitation was projected to increase by 43.250 mm (σ = 88.818 mm) over the same time period. On average, GO predictions increased (*µ*_Δ*GO*_ = 2.113) from 1990 to 2090 across all populations, but there was meaningful variation in ΔGO among populations (σ_Δ*GO*_ = 5.747). While 23 populations (54.8%) experienced an increase in GO between 1990 and 2090, nineteen populations (45.2%) experienced a decrease in GO over the same time period. Thus, our projections suggest that approximately half of the *Lycaeides* populations will experience greater maladaptation to climatic conditions over the course of the century (1990-2090).

**Figure 1:**
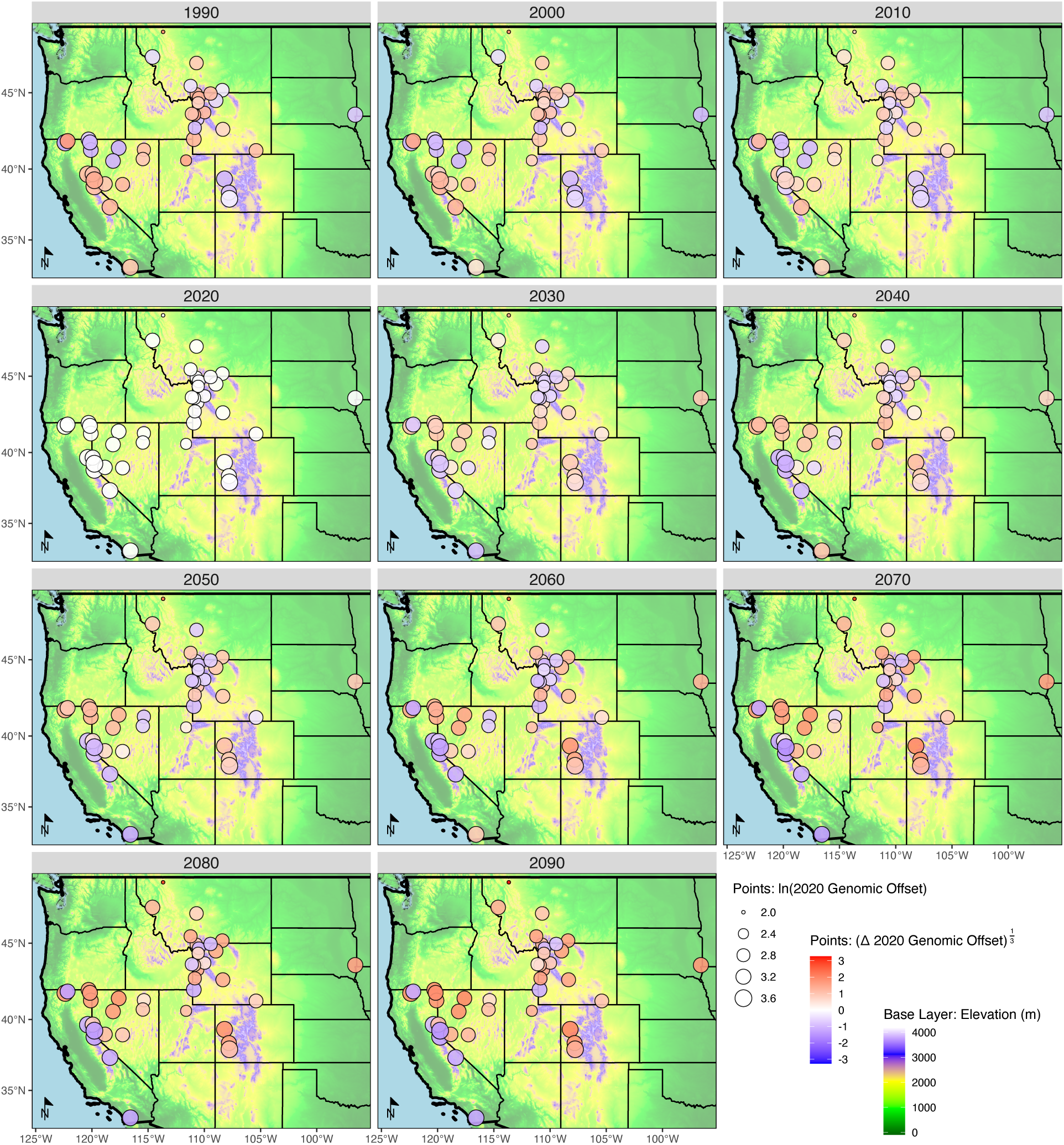
Spatial distribution of *Lycaeides* populations with genomic offset (GO) projections. Point size represents contemporary GO, while point color represents the change in GO since 2020 (i.e., decadal GO minus 2020 GO). Blue points represent lower malaptation compared to contemporary conditions, whereas red points represent higher maladptation compared to contemporary conditions. Elevation base layer from the North America Elevation 1 Kilometer Resolution GRID.

**Figure 2:**
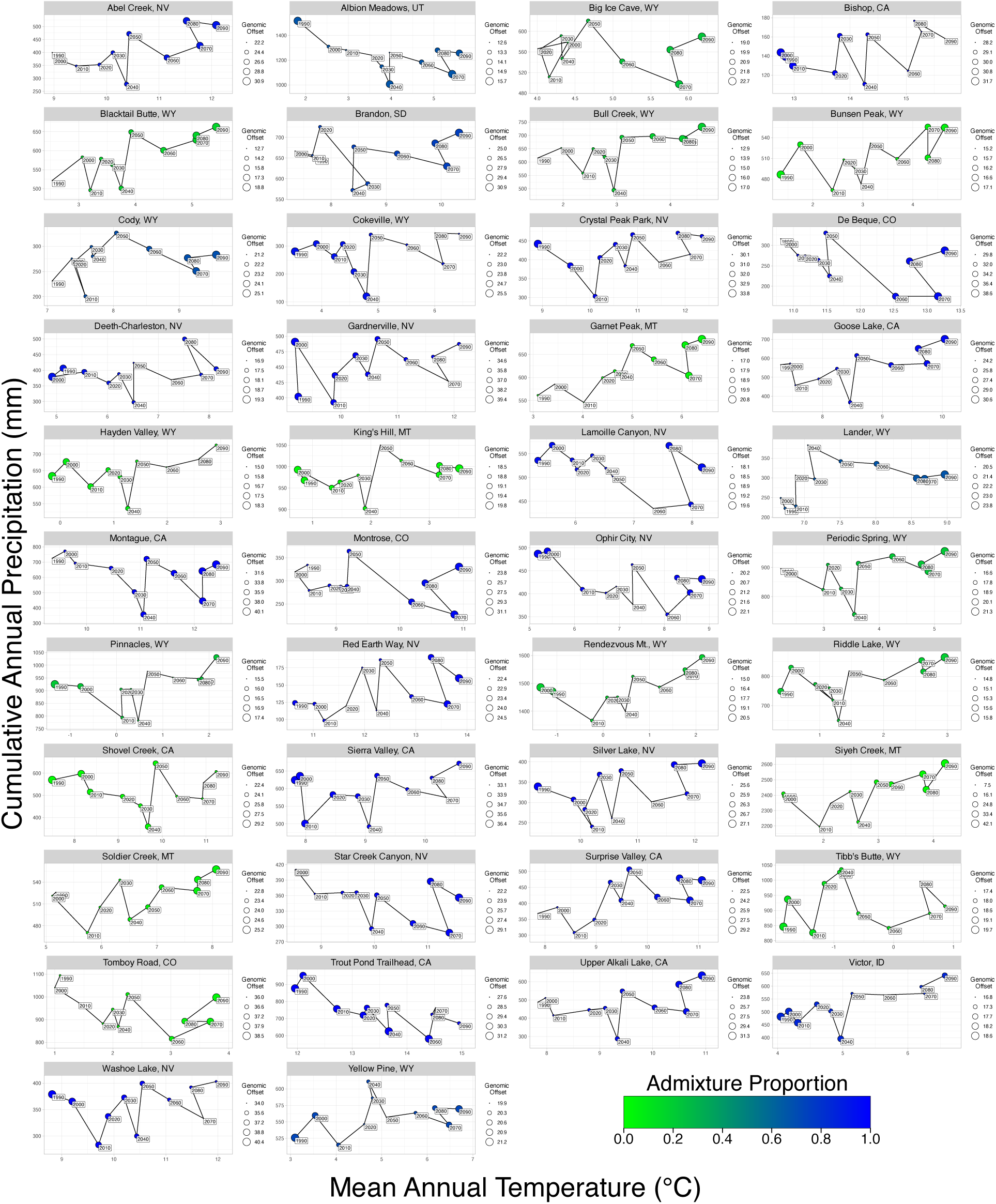
Climatic conditions of *Lycaeides* populations with genomic offset (GO) projections. Point size represents GO projections for each decade, and point color represents a population’s admixture proportion. An admixture proportion of one represents *Lycaeides melissa*, whereas an admixture proportion of zero represents *Lycaeides idas*.

The linear mixed-effects model of GO through time is shown in Figure 3. We found a statistically-significant (𝛼 = 0.05) positive relationship between log-transformed GO and year (𝛽 = 9.930 × 10*^−^*^4^, *p*-value *<* 0.001). Given the logarithmic scale of the genomic offset response, these results suggest an average increase in GO of 1.0% per decade or 10.4% per century. The linear mixed-effects model also found meaningful variation in GO between populations, with the standard deviation of random effects exceeding the residual standard deviation (σ*_population_* = 0.283, σ*_residual_* = 0.107). Taken together, the ΔGO calculations and linear mixed-effects model both suggest an average increase in GO between 1990 and 2090, with substantial variation in GO between populations.

**Figure 3:**
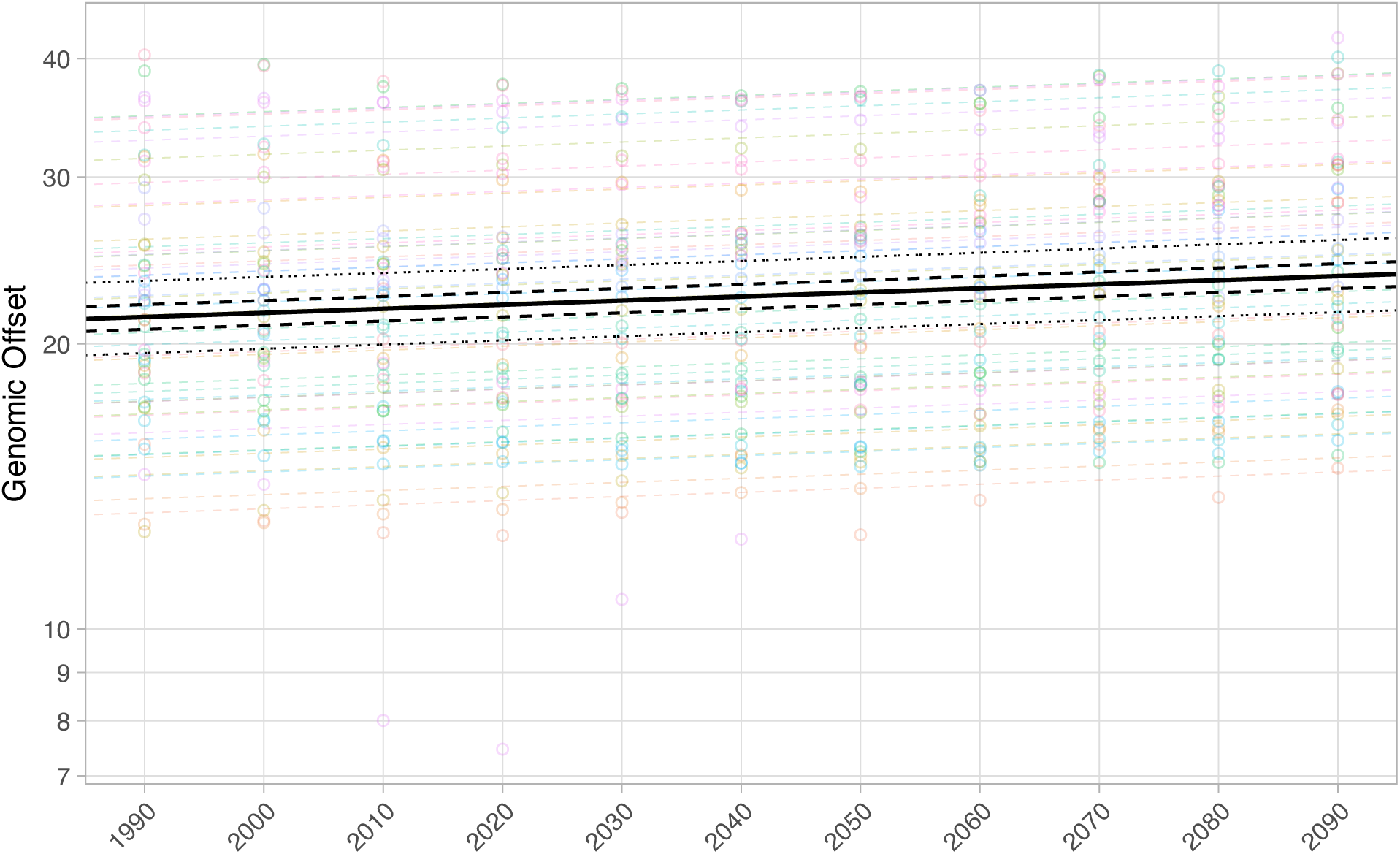
Linear mixed-effects model of genomic offset projections through time. The colored dashed lines represent population predictions. The black solid line represents expected values, the black dashed lines represent 50% confidence limits, and the black dotted lines represent 95% confidence limits. Note the logarithmic scale on the y-axis.

### Gene enrichment by environmental association

Gene enrichment tests for the best-fit model predictors (Table S4) found statistically significant gene associations for admixture, temperature, and the two-way interactions with admixture (i.e., temperature and precipitation). However, the magnitude of gene enrichment was weak, with all significant tests exhibiting enrichments ≤ 1.7% higher than the overall proportion of SNPs on or near genes (84.7%; Table S4). As enrichment results are dependent on the accuracy of functional annotations, we opted to focus on the relative (as opposed to absolute) strength of gene associations between predictors. The strongest enrichments occurred for admixture, temperature, and their interaction (proportion ratios of 1.014, 1.012, and 1.017, respectively; *p*-values *<* 0.001), whereas the only statistically-significant enrichment involving precipitation was for its interaction with admixture (proportion ratio = 1.016, *p*-value = 0.017). Meanwhile, precipitation (proportion ratio = 0.999, *p*-value = 0.587) and the three-way interaction (proportion ratio = 1.003, *p*-value = 0.365) produced non-significant enrichment results (Table S4). Taken together, gene enrichments were strongest for admixture and temperature effects, while gene enrichments were weakest for effects involving precipitation.

Gene ontology enrichment tests for the best-fit model predictors produced 15 statistically-significant predictor-ontology combinations (Table S5). The greatest enrichment magnitude was expressed by two gene ontologies associated with the three-way interaction. SNPs annotated with gene ontology numbers #0000289 (biological process: nuclear-transcribed mRNA poly[A] tail shortening) and #0031251 (cellular component: PAN complex) were both 19.6 times more common (as a proportion ratio) in the pool of SNPs significantly associated with the three-way interaction than compared to all SNPs in the study (FDR-adjusted *p*-values *<* 0.001). Two gene ontologies exhibited significant enrichment across multiple predictors. Gene ontology number #1990316 (cellular component: Atg1/ULK1 kinase complex) was significantly associated with temperature (proportion ratio = 6.00, FDR-adjusted *p*-value *<* 0.001) and the admixture-temperature interaction (proportion ratio = 7.718, FDR-adjusted *p*-value *<* 0.001). Gene ontology number #0004563 (molecular function: 𝛽-N-acetylhexosaminidase activity) was significantly associated with precipitation (proportion ratio = 7.813, FDR-adjusted *p*-value *<* 0.001) and the three-way interaction (proportion ratio = 6.302, FDR-adjusted *p*-value *<* 0.001).

### Relationship between GOs and demographic factors

As identified by LOO-CV elppd, the best-fit multiple linear regression model between 2020 GO and demographic parameters included effective migration and effective diversity as predictors (elppd = *−*14.777, *R*^2^ = 0.273, *w_i_* = 0.360). The model with the second-highest elppd included only effective diversity as a predictor (elppd = *−*15.504, *R*^2^ = 0.216, *w_i_* = 0.174). No other competing models were within two units of the best-fitting model by *−*2 × elppd (Table 2). Within the best-fit model, effective diversity (partial *R*^2^ = 0.225) and effective migration (partial *R*^2^ = 0.073) both explained a non-trivial amount of variation in contemporary GO. Both predictors exhibited either statistically-significant (𝛼 = 0.05) or marginally-significant (𝛼 = 0.1) negative relationships with genomic offset. For effective diversity (𝛽 = *−* 0.154, *p*-value = 0.002; Figure 4A), this relationship is consistent with our hypothesis that genetic diversity provides adaptive capacity. For effective migration (𝛽 = *−* 0.080, *p*-value = 0.088; Figure 4B), this relationship is consistent with our hypothesis that gene flow contributes potentially-adaptive alleles. By sum of weights (sw; Burnham & Anderson, 2004), effective diversity (sw = 0.711) was the most important predictor of genomic offset, while effective migration (sw = 0.660) was a slightly weaker predictor. The importance of *N_e_* in predicting genomic offset (sw = 0.255) was substantially lower than the other demographic parameters.

**Figure 4:**
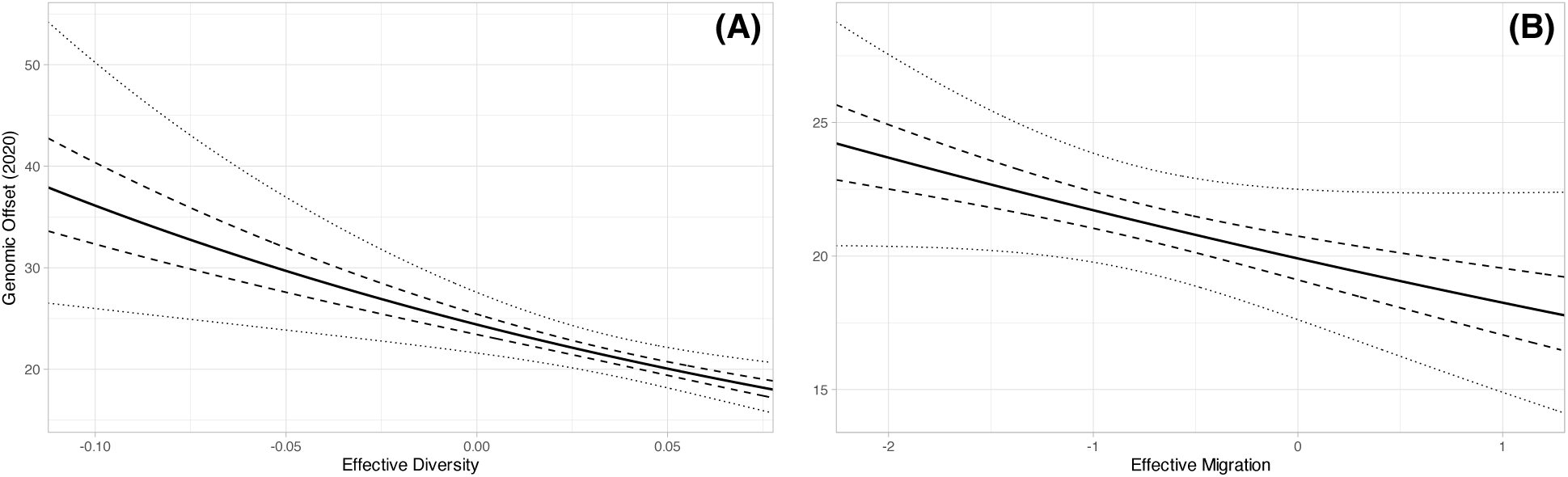
Back-transformed marginal effects of effective diversity (A) and effective migration (B) on contemporary genomic offset (GO). Predictions generated from the best-fit multiple linear regression model of demographic parameters on 2020 GO while holding other predictors constant at their mean values (*x̄_migration_* = *−*0.805, *x̄_diversity_* = 0.034). The x-axes span the range of effective diversity and effective migration values present in the population data. The solid lines represent expected values, the dashed lines represent 50% confidence limits, and the dotted lines represent 95% confidence limits.

**Table 2:**
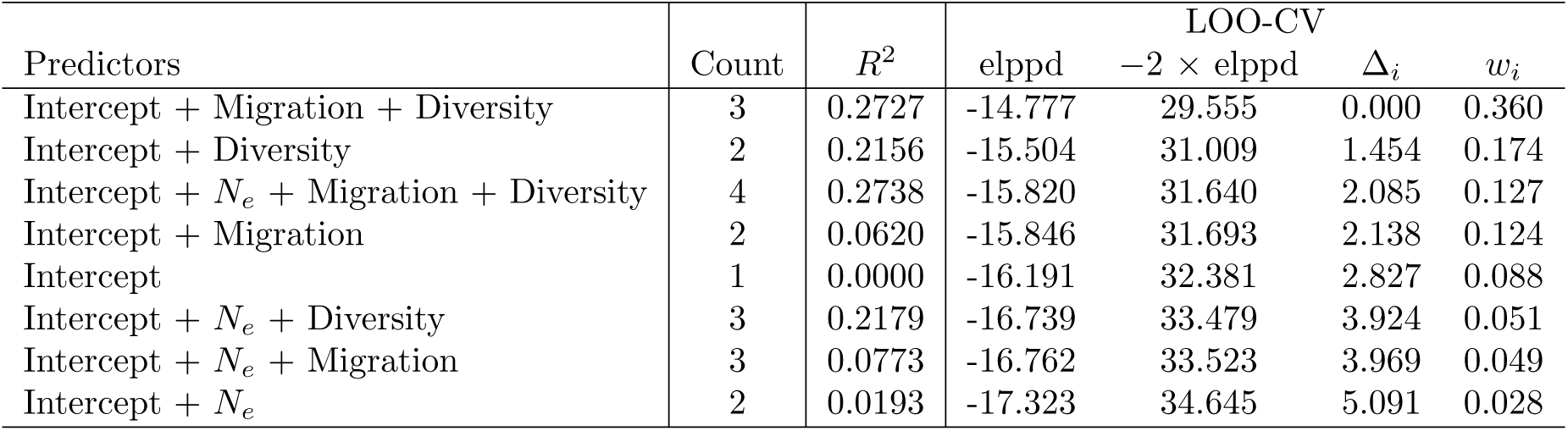
Summary of competing demographic models. Predictor combinations are displayed for each linear regression on contemporary genomic offset. The count field contains the number of model predictors. The best-fit model (first record) has the highest leave-one-out cross-validated (LOO-CV) expected log pointwise predictive density (elppd), which estimates out-of-sample predictive performance. Δ*_i_* is the difference in *−*2 × elppd compared to the best-fit model. *w_i_* represents model weights. Models are sorted by increasing Δ*_i_*.

## Discussion

Predicting which populations or species are most likely to face an elevated risk of decline or extinction due to anthropogenic climate change is an urgent goal for both biologists and society (Bernatchez *et al*., 2024). Consequently, the potential to enhance predictive power by incorporating genomic data has generated considerable interest in the scientific community (Bay *et al*., 2018b; Capblancq *et al*., 2020; Exposito-Alonso *et al*., 2022), even leading to real-world applications of genomic forecasting (Beck *et al*., 2025; Jacquemart *et al*., 2025). However, recent work has shown that the utility of genomic forecasts can be system- and application-specific, that they complement rather than replace alternative methods, and that substantial validation work remains to be done (e.g., Lotterhos, 2024; Rellstab & Keller, 2025; Fitzpatrick *et al*., 2026). Our study contributes to this ongoing debate in several key ways. First, we extend the focus to insects, specifically butterflies, which are known to be experiencing substantial declines (Halsch *et al*., 2021; Harvey *et al*., 2023; Forister *et al*., 2024; Edwards *et al*., 2025). Our study system, *Lycaeides*, includes endangered and recently extinct taxa (Forister *et al*., 2011; Arnold, 1980, 1993). Second, we adopt population genetic estimates of demographic correlates to explain and validate the GO estimates. This approach is important because these metrics require no data beyond that used to generate GOs, making them broadly applicable to non-model systems. Finally, we examine GOs across a broad spatial and temporal scale, highlighting the heterogeneity in predicted risk of maladaptation. Below, we first discuss the relevance of our results for understanding climate adaptation and then turn to the issue of the utility of GOs.

### Genetics of climate adaptation

We found that admixture proportion, mean annual temperature, and cumulative annual precipitation explained a substantial amount of variation in allele frequencies across *Lycaeides* populations (best-fit GEA *R*^2^ = 47.6%). While admixture proportion was the single most important predictor (Table S3), the climatic predictors also contributed meaningful explanatory power. By generalizing *R*^2^ for model comparison, we found that, after controlling for the effect of admixture proportion, the climatic predictors and their interactions explained 22.5% of the remaining variation in population allele frequencies. This high explanatory power of climatic predictors is consistent with a polygenic architecture of climate adaptation. Within the best-fit GEA model, partial *R*^2^ statistics suggested that admixture, temperature, and their interaction were most important in explaining genetic variation, whereas precipitation and its interactions contributed less (Table S3). Thus, a surprising amount of genetic variation can be explained by the climatic predictors, particularly temperature, and their interactions with admixture suggest that climatic adaptation is dependent on genetic background. We further found that alleles associated with climatic adaptation were unevenly distributed across the genome, with climate-associated SNPs concentrated on chromosomes Z and 1. These results are consistent with documented effects of temperature and precipitation on butterfly abundance and phenology, and with the fact that such effects often depend on genetic background (i.e., vary among populations and species) (Nice *et al*., 2019; Forister *et al*., 2023; Reis *et al*., 2026). Our results are also consistent with many traits in butterflies being strongly influenced by genetic variation on the Z chromosome (Sperling, 1994; Mongue *et al*., 2022; Gompert *et al*., 2025b).

We found significant gene enrichment for most GEA predictors (Table S4), suggesting the potential for locally-adapted functional traits. Functional annotations identified 13 gene ontologies that were significantly associated with GEA predictors, of which 10 were significantly associated with climatic predictors or their interactions (Table S5). Two gene ontologies were associated with multiple predictors: the Atg1/ULK1 kinase complex (temperature and admixture-temperature interaction) and 𝛽-N-acetylhexosaminidase activity (precipitation and three-way interaction). The Atg1/ULK1 kinase complex is involved in autophagy, which can occur in response to starvation. The insect glycoside hydrolase family 20 𝛽-N-acetylhexosaminidases (HEXs) are known to be key enzymes for chitin degradation, which plays a central role in molting (Zhang *et al*., 2022). Based on these associations, *Lycaeides* populations may exhibit temperature-related adaptations to starvation and precipitation-related adaptations to molting. Nonetheless, much remains unknown about the genetic basis of climate adaptation in this system, including the potential importance of adaptive changes in gene regulation and plasticity, and the role of chromosomal rearrangements (i.e., structural variation), both of which have increasingly been shown to be important for local adaptation (e.g., Todesco *et al*., 2020; Oomen & Hutchings, 2022; Nosil *et al*., 2023; Battlay *et al*., 2025; Gompert *et al*., 2025a).

### Genomic forecasting of maladaptation and extinction risk

Although temporal trends in GO varied substantially among populations – increasing for some (e.g., MON and SYC) and decreasing for others (e.g., BHP and TPT) – we projected an average increase in GO of 10.4% from 1990 to 2090 (Figure 3). These results are consistent with the hypothesis that the risk of maladaptation and extinction due to climate change will increase over time, as suggested more generally by a number of studies (e.g., Thomas *et al*., 2004; Urban, 2015). However, this genomic forecast should be interpreted with considerable caution. Extrapolating GEA relationships to the novel climate conditions that are expected to prevail in the future is inherently risky (Fitzpatrick & Hargrove, 2009; Rellstab & Keller, 2025). Moreover, adaptation to climate could keep pace with, or only slightly lag behind, climate change, such that the difference between the actual and (near)optimal genetic composition might not change substantially over time (Colautti & Barrett, 2013; Hoffmann, 2017; Rudman *et al*., 2022), whereas our GOs reflect how the current population might perform under future climatic conditions.

Although we hypothesized a negative relationship between GO and contemporary effective population size, *N_e_* was not included in the best-fit regression model between contemporary GO and demographic parameters. Instead, we found *N_e_* to be the weakest predictor of contemporary GO by sum of weights. For the other demographic parameters, we found either statistically-significant or marginally-significant negative relationships between contemporary GO and the best-fit model predictors of effective diversity and effective migration (Figures 4A and 4B, respectively). By partial *R*^2^, effective diversity was the most important predictor of contemporary GO, with the combined effects of effective diversity and effective migration explaining over a quarter of the variation in contemporary GO (*R*^2^ = 27.3%).

As potential explanations, populations with high genetic diversity may have greater adaptive capacity, which in turn may reduce their GOs (Ørsted *et al*., 2019; Kardos *et al*., 2021; Aitken *et al*., 2024). Similarly, high migration rates may provide an influx of genes with adaptive potential, thereby enabling populations to maintain local adaptation despite changing environmental conditions (Nicotra *et al*., 2015; Torda & Quigley, 2022; Aitken *et al*., 2024). In contrast, current (effective) population sizes, or even recent demographic changes, might not yet be reflected in levels of maladaptation (i.e., there might be a drift debt; Pinto *et al*., 2024; Gargiulo *et al*., 2025), but may have stronger effects on a population’s ability to keep evolutionary pace with climate change over longer timescales. In terms of gene flow and connectivity, admixture and introgressive hybridization are increasingly being recognized as potential sources of adaptive variation that allow populations to adapt to novel or changing environments (Taylor *et al*., 2015; Brauer *et al*., 2023; Hord *et al*., 2025). This is particularly relevant for *Lycaeides* butterflies given the prevalence of admixture within this species complex (Gompert *et al*., 2010, 2014; Chaturvedi *et al*., 2020; Gompert *et al*., 2025b), as well as the substantial explanatory power of past admixture for patterns of allele frequency variation documented in this paper.

Importantly, here we focused on a single method for genomic forecasts of past, present, and future maladaptation and on a single set of population genetic estimates of demographic parameters to explain and validate these predictions. Other methods, especially those that allow for nonlinear relationships between climatic variables and genetic variation, could produce qualitatively distinct predictions of maladaptation and extinction risk (Ferrier *et al*., 2007; Fitzpatrick & Keller, 2015; Archambeau *et al*., 2026). Different validation approaches could also yield substantially different results (Lotterhos, 2024; Rellstab & Keller, 2025). A key strength of our approach is that it does not rely on additional data, but it also does not directly measure fitness or extinction risk. Laboratory (growth chamber) measurements of fitness components or correlates under standardized conditions (i.e., “common gardens”) are practical for insects, including *Lycaeides* (Forister *et al*., 2009; Gompert *et al*., 2019, 2022), but it is unclear how well these measures relate to fitness and population persistence in nature (Forister *et al*., 2020). Perhaps the most relevant approach for validating extinction risk is to test predictions against demographic monitoring data, as has been done for some bird species (e.g., Bay *et al*., 2018b). Fortunately, such monitoring data also exist for butterflies (Forister *et al*., 2021), and we have future work underway to validate genomic predictions against these data in *Lycaeides* butterflies (Reis *et al*., manuscript in preparation). Only through such extensive validation will we ultimately come to understand when and in what ways genomic forecasting can guide conservation efforts versus mislead them.

## Data archiving

### Data availability

Raw sequence data are available in the NCBI Sequence Read Archives associated with Gompert *et al*. (2014) (BioProject ID: PRJNA246037) and Chaturvedi *et al*. (2018) (BioProject ID: PRJNA432816). Population metadata are included in Table 1.

### Code availability

Code used in data processing and analyses are available on GitHub: https://github.com/Urodelan/LycaeidesGenomicOffset; https://github.com/zgompert/LycaeidesGenomicOffsetGenetics; https://github.com/chaturvedi-lab/Lmelissa_genome_annotation.

## Author contributions

K.B.G. performed genotype-environment-association analyses, genomic offset predictions, and enrichment tests. S.C. and Z.G. performed sequence alignments. S.C. performed genome and functional annotations. Z.G. estimated demographic parameters. S.C., L.K.L., and Z.G. generated the data. K.B.G, S.C., and Z.G. wrote the manuscript. All authors contributed to manuscript revisions.

## Research funding

This research was financially supported by Utah State University and the National Science Foundation (NSF DEB 1844941 to ZG).

