## Supplement for "Genomic forecasts of maladptation in *Lycaeides* butterflies"

### Supplemental Text

#### Genome annotation

We annotated structural and functional features of the new *L. melissa* genome using BRAKER3 (Gabriel *et al.*, 2024), an evidence-based annotation pipeline that integrates repeat masking, RNA-seq data, and protein evidence for whole-genome gene prediction. Prior to annotation, we constructed a *de novo* repeat library for the *L. melissa* genome using RepeatModeler2 (Flynn *et al.*, 2020) with default settings. Repetitive elements were then soft-masked with RepeatMasker v4.0.6 (Smit, 2004), including additional long terminal repeat identification with the `-LTRStruct` option.

To provide transcript evidence for BRAKER3, we used RNA-seq data from 24 *L. melissa* transcriptomes. Raw paired-end reads were first assessed with FastQC v0.11.7 (Andrews 2010). Adapters and low-quality bases were removed using Trim Galore v2.6.6 (Krueger, 2015), and reads shorter than 50 bp after trimming were discarded. The filtered paired-end reads were then aligned to the *L. melissa* reference genome using STAR v2.7.11 (Dobin *et al.*, 2013). For this, we first generated a genome index with STAR `--genomeGenerate` and then mapped reads to the genome to produce sorted BAM files for downstream annotation in BRAKER3.

BRAKER3 was run using the soft-masked genome assembly together with the RNA-seq alignments and protein evidence to predict gene models. Functional annotation of predicted proteins was subsequently performed using BLASTP against the UniProt/Swiss-Prot database and InterProScan v5.32-71.0 to assign protein domains and Gene Ontology terms (Jones *et al.*, 2014). The final annotation included 11,247 putative genes, 48,765 putative coding sequences, and 8893 UTR sequences.

### Supplemental Tables

Table S1: Estimates of contemporary effective population size ($N_e$) based on patterns of linkage disequilibrium among unlinked loci. Point estimates and the 25th and 75th percentiles (%) from resampling subsets of locus pairs (100 pairs of 10,000 SNPs) are shown for each population. See Table 1 for detailed population information.

| Population ID | $N_e$ | 25th% | 75th% |
|---|---|---|---|
| ABC | 319.03 | 193.00 | 572.15 |
| ABM | 403.84 | 323.06 | 570.91 |
| BCR | 161.53 | 148.92 | 183.95 |
| BHP | 14.45 | 13.75 | 15.25 |
| BIC | 254.77 | 165.57 | 497.57 |
| BNP | 267.41 | 166.08 | 514.89 |
| BSD | 40.93 | 37.87 | 44.00 |
| BTB | 728.90 | 564.81 | 1315.21 |
| CDY | 134.14 | 111.98 | 177.40 |
| CKV | 39.10 | 33.40 | 45.34 |
| DBQ | 142.26 | 106.70 | 191.81 |
| DCR | 105.01 | 86.55 | 122.66 |
| GLA | 60.03 | 54.41 | 64.16 |
| GNP | 117.45 | 102.37 | 164.13 |
| GVL | 58.59 | 51.47 | 66.08 |
| HNV | 268.98 | 168.07 | 445.46 |
| KHL | 176.82 | 124.02 | 289.04 |
| LAN | 337.17 | 252.22 | 558.22 |
| LCA | 58.29 | 52.56 | 65.93 |
| MON | 423.37 | 302.59 | 848.58 |
| MTU | 472.28 | 297.68 | 725.74 |
| OCY | 90.64 | 76.50 | 104.76 |
| PIN | 236.58 | 184.54 | 405.57 |
| PSP | 78.76 | 68.21 | 90.98 |
| RDL | 54.55 | 50.93 | 58.66 |
| REW | 124.95 | 102.94 | 156.79 |
| RNV | 153.16 | 131.31 | 177.52 |
| SCC | 111.93 | 91.35 | 155.73 |
| SDC | 172.93 | 138.10 | 259.25 |
| SHC | 62.12 | 55.64 | 70.65 |
| SLA | 211.48 | 152.76 | 411.07 |
| SUV | 456.90 | 244.71 | 1167.99 |
| SVY | 19.48 | 18.91 | 20.44 |
| SYC | 101.80 | 86.41 | 124.56 |
| TBY | 101.19 | 86.89 | 117.13 |
| TIB | 80.05 | 69.12 | 90.29 |
| TPT | 186.91 | 144.18 | 371.17 |
| UAL | 462.58 | 264.23 | 704.41 |
| VCP | 242.64 | 174.71 | 338.71 |
| VIC | 229.78 | 172.06 | 397.57 |
| WAL | 124.88 | 102.24 | 168.70 |
| YWP | 97.94 | 82.12 | 127.54 |

Table S2: Summary of competing genotype-environment-association models. Predictor combinations are displayed for each multivariate-normal linear regression model. The count field contains the number of model predictors. Models are sorted by decreasing adjusted $R^2$.

| Predictors | Count | $R^2$ | Adjusted $R^2$ |
|---|---|---|---|
| Intercept + Admixture + Temperature + Precipitation + Admixture:Temperature + Admixture:Precipitation + Admixture:Temperature:Precipitation | 7 | 0.4763 | 0.3865 |
| Intercept + Admixture + Temperature + Precipitation + Admixture:Temperature + Temperature:Precipitation + Admixture:Temperature:Precipitation | 7 | 0.4759 | 0.3860 |
| Intercept + Admixture + Temperature + Admixture:Temperature + Admixture:Precipitation + Admixture:Temperature:Precipitation | 6 | 0.4601 | 0.3851 |
| Intercept + Admixture + Temperature + Admixture:Temperature + Temperature:Precipitation + Admixture:Temperature:Precipitation | 6 | 0.4588 | 0.3836 |
| Intercept + Admixture + Temperature + Admixture:Temperature + Admixture:Precipitation + Temperature:Precipitation + Admixture:Temperature:Precipitation | 7 | 0.4736 | 0.3833 |
| Intercept + Admixture + Temperature + Precipitation + Admixture:Temperature + Admixture:Precipitation + Temperature:Precipitation + Admixture:Temperature:Precipitation | 8 | 0.4883 | 0.3830 |
| Intercept + Admixture + Temperature + Precipitation + Admixture:Temperature + Admixture:Precipitation + Temperature:Precipitation | 7 | 0.4732 | 0.3829 |
| Intercept + Admixture + Temperature + Precipitation + Admixture:Temperature + Temperature:Precipitation | 6 | 0.4569 | 0.3815 |
| Intercept + Admixture + Temperature + Precipitation + Admixture:Temperature + Admixture:Precipitation | 6 | 0.4567 | 0.3812 |
| Intercept + Admixture + Temperature + Precipitation + Admixture:Temperature + Admixture:Temperature:Precipitation | 6 | 0.4536 | 0.3777 |
| Intercept + Admixture + Temperature + Admixture:Temperature | 4 | 0.4229 | 0.3773 |
| Intercept + Admixture + Temperature + Admixture:Temperature + Admixture:Precipitation + Temperature:Precipitation | 6 | 0.4528 | 0.3768 |
| Intercept + Admixture + Temperature + Admixture:Temperature + Admixture:Temperature:Precipitation | 5 | 0.4372 | 0.3764 |
| Intercept + Admixture + Temperature + Admixture:Temperature + Admixture:Precipitation | 5 | 0.4370 | 0.3762 |
| Intercept + Admixture + Temperature + Admixture:Temperature + Temperature:Precipitation | 5 | 0.4368 | 0.3759 |
| Intercept + Admixture + Temperature + Precipitation + Admixture:Temperature | 5 | 0.4355 | 0.3744 |
| Intercept + Admixture + Temperature + Precipitation + Admixture:Precipitation + Admixture:Temperature:Precipitation | 6 | 0.4350 | 0.3565 |
| Intercept + Admixture + Temperature + Precipitation + Temperature:Precipitation + Admixture:Temperature:Precipitation | 6 | 0.4341 | 0.3555 |
| Intercept + Admixture + Temperature + Temperature:Precipitation + Admixture:Temperature:Precipitation | 5 | 0.4166 | 0.3535 |
| Intercept + Admixture + Temperature + Admixture:Precipitation + Admixture:Temperature:Precipitation | 5 | 0.4154 | 0.3522 |
| Intercept + Admixture + Temperature + Admixture:Precipitation + Temperature:Precipitation + Admixture:Temperature:Precipitation | 6 | 0.4312 | 0.3522 |
| Intercept + Admixture + Temperature + Precipitation + Admixture:Precipitation + Temperature:Precipitation + Admixture:Temperature:Precipitation | 7 | 0.4465 | 0.3517 |
| Intercept + Admixture + Temperature + Precipitation + Admixture:Precipitation + Temperature:Precipitation | 6 | 0.4300 | 0.3508 |
| Intercept + Admixture + Temperature + Precipitation + Temperature:Precipitation | 5 | 0.4130 | 0.3495 |
| Intercept + Admixture + Temperature + Admixture:Precipitation + Temperature:Precipitation | 5 | 0.4107 | 0.3470 |
| Intercept + Admixture + Temperature + Precipitation + Admixture:Precipitation | 5 | 0.4106 | 0.3469 |
| Intercept + Admixture + Temperature + Precipitation + Admixture:Temperature:Precipitation | 5 | 0.4105 | 0.3468 |
| Intercept + Admixture + Temperature + Temperature:Precipitation | 4 | 0.3940 | 0.3462 |
| Intercept + Admixture + Temperature | 3 | 0.3775 | 0.3455 |
| Intercept + Admixture + Precipitation + Admixture:Temperature + Temperature:Precipitation + Admixture:Temperature:Precipitation | 6 | 0.4243 | 0.3444 |
| Intercept + Admixture + Temperature + Precipitation | 4 | 0.3921 | 0.3441 |
| Intercept + Admixture + Temperature + Admixture:Precipitation | 4 | 0.3916 | 0.3436 |
| Intercept + Admixture + Precipitation + Admixture:Temperature + Admixture:Precipitation + Admixture:Temperature:Precipitation | 6 | 0.4235 | 0.3434 |
| Intercept + Admixture + Admixture:Temperature + Admixture:Precipitation + Admixture:Temperature:Precipitation | 5 | 0.4072 | 0.3431 |
| Intercept + Admixture + Admixture:Temperature + Temperature:Precipitation + Admixture:Temperature:Precipitation | 5 | 0.4070 | 0.3429 |
| Intercept + Admixture + Temperature + Admixture:Temperature:Precipitation | 4 | 0.3909 | 0.3428 |
| Intercept + Admixture + Precipitation + Admixture:Temperature + Admixture:Precipitation + Temperature:Precipitation | 6 | 0.4218 | 0.3415 |
| Intercept + Admixture + Precipitation + Admixture:Temperature + Temperature:Precipitation | 5 | 0.4055 | 0.3413 |
| Intercept + Admixture + Admixture:Temperature + Admixture:Precipitation + Temperature:Precipitation + Admixture:Temperature:Precipitation | 6 | 0.4207 | 0.3403 |
| Intercept + Admixture + Precipitation + Admixture:Temperature + Admixture:Precipitation | 5 | 0.4045 | 0.3401 |
| Intercept + Admixture + Precipitation + Admixture:Temperature + Admixture:Precipitation + Temperature:Precipitation + Admixture:Temperature:Precipitation | 7 | 0.4357 | 0.3389 |
| Intercept + Admixture + Admixture:Temperature | 3 | 0.3699 | 0.3376 |
| Intercept + Admixture + Precipitation + Admixture:Temperature + Admixture:Temperature:Precipitation | 5 | 0.4016 | 0.3369 |
| Intercept + Admixture + Admixture:Temperature + Admixture:Temperature:Precipitation | 4 | 0.3851 | 0.3365 |
| Intercept + Admixture + Admixture:Temperature + Admixture:Precipitation + Temperature:Precipitation | 5 | 0.4008 | 0.3361 |
| Intercept + Admixture + Admixture:Temperature + Admixture:Precipitation | 4 | 0.3841 | 0.3355 |
| Intercept + Admixture + Admixture:Temperature + Temperature:Precipitation | 4 | 0.3836 | 0.3350 |
| Intercept + Admixture + Precipitation + Admixture:Temperature | 4 | 0.3836 | 0.3349 |
| Intercept + Admixture + Precipitation + Temperature:Precipitation + Admixture:Temperature:Precipitation | 5 | 0.3835 | 0.3169 |
| Intercept + Admixture + Precipitation + Admixture:Precipitation + Admixture:Temperature:Precipitation | 5 | 0.3831 | 0.3164 |
| Intercept + Admixture + Temperature:Precipitation + Admixture:Temperature:Precipitation | 4 | 0.3657 | 0.3156 |
| Intercept + Admixture + Admixture:Precipitation + Admixture:Temperature:Precipitation | 4 | 0.3636 | 0.3134 |
| Intercept + Admixture + Admixture:Precipitation + Temperature:Precipitation + Admixture:Temperature:Precipitation | 5 | 0.3794 | 0.3123 |
| Intercept + Admixture + Precipitation + Admixture:Precipitation + Temperature:Precipitation | 5 | 0.3791 | 0.3120 |
| Intercept + Admixture + Precipitation + Temperature:Precipitation | 4 | 0.3621 | 0.3118 |
| Intercept + Admixture + Precipitation + Admixture:Precipitation + Temperature:Precipitation + Admixture:Temperature:Precipitation | 6 | 0.3949 | 0.3109 |
| Intercept + Admixture + Admixture:Precipitation + Temperature:Precipitation | 4 | 0.3586 | 0.3079 |
| Intercept + Admixture | 2 | 0.3245 | 0.3076 |
| Intercept + Admixture + Precipitation + Admixture:Precipitation | 4 | 0.3576 | 0.3068 |
| Intercept + Admixture + Temperature:Precipitation | 3 | 0.3402 | 0.3064 |
| Intercept + Admixture + Precipitation + Admixture:Temperature:Precipitation | 4 | 0.3562 | 0.3054 |
| Intercept + Admixture + Precipitation | 3 | 0.3376 | 0.3036 |
| Intercept + Admixture + Admixture:Precipitation | 3 | 0.3374 | 0.3034 |
| Intercept + Admixture + Admixture:Temperature:Precipitation | 3 | 0.3368 | 0.3027 |
| Intercept + Temperature + Admixture:Temperature + Admixture:Precipitation + Temperature:Precipitation + Admixture:Temperature:Precipitation | 6 | 0.3248 | 0.2310 |
| Intercept + Temperature + Admixture:Temperature + Admixture:Precipitation + Admixture:Temperature:Precipitation | 5 | 0.3048 | 0.2296 |
| Intercept + Temperature + Precipitation + Admixture:Temperature + Admixture:Precipitation + Admixture:Temperature:Precipitation | 6 | 0.3212 | 0.2269 |
| Intercept + Temperature + Precipitation + Admixture:Temperature + Admixture:Precipitation + Temperature:Precipitation + Admixture:Temperature:Precipitation | 7 | 0.3393 | 0.2261 |
| Intercept + Temperature + Precipitation + Admixture:Temperature + Admixture:Precipitation | 5 | 0.2988 | 0.2230 |
| Intercept + Temperature + Precipitation + Admixture:Temperature + Admixture:Precipitation + Temperature:Precipitation | 6 | 0.3178 | 0.2230 |
| Intercept + Temperature + Admixture:Temperature + Temperature:Precipitation + Admixture:Temperature:Precipitation | 5 | 0.2981 | 0.2222 |
| Intercept + Temperature + Precipitation + Admixture:Temperature + Temperature:Precipitation | 5 | 0.2968 | 0.2208 |
| Intercept + Temperature + Precipitation + Admixture:Temperature + Temperature:Precipitation + Admixture:Temperature:Precipitation | 6 | 0.3156 | 0.2205 |
| Intercept + Temperature + Admixture:Temperature | 3 | 0.2581 | 0.2200 |
| Intercept + Temperature + Admixture:Temperature + Admixture:Temperature:Precipitation | 4 | 0.2766 | 0.2194 |
| Intercept + Temperature + Precipitation + Admixture:Temperature | 4 | 0.2756 | 0.2184 |
| Intercept + Temperature + Precipitation + Admixture:Temperature + Admixture:Temperature:Precipitation | 5 | 0.2934 | 0.2170 |
| Intercept + Temperature + Admixture:Temperature + Temperature:Precipitation | 4 | 0.2731 | 0.2158 |
| Intercept + Temperature + Admixture:Temperature + Admixture:Precipitation + Temperature:Precipitation | 5 | 0.2910 | 0.2144 |
| Intercept + Temperature + Admixture:Temperature + Admixture:Precipitation | 4 | 0.2718 | 0.2143 |
| Intercept + Temperature + Admixture:Precipitation + Temperature:Precipitation + Admixture:Temperature:Precipitation | 5 | 0.2830 | 0.2055 |
| Intercept + Temperature + Precipitation + Admixture:Precipitation + Temperature:Precipitation + Admixture:Temperature:Precipitation | 6 | 0.2982 | 0.2007 |
| Intercept + Temperature + Precipitation + Admixture:Precipitation + Admixture:Temperature:Precipitation | 5 | 0.2784 | 0.2004 |
| Intercept + Temperature + Admixture:Precipitation + Admixture:Temperature:Precipitation | 4 | 0.2578 | 0.1993 |
| Intercept + Temperature + Precipitation + Admixture:Precipitation + Temperature:Precipitation | 5 | 0.2758 | 0.1975 |
| Intercept + Temperature + Temperature:Precipitation + Admixture:Temperature:Precipitation | 4 | 0.2561 | 0.1973 |
| Intercept + Temperature + Precipitation + Temperature:Precipitation + Admixture:Temperature:Precipitation | 5 | 0.2744 | 0.1959 |
| Intercept + Temperature + Precipitation + Temperature:Precipitation | 4 | 0.2536 | 0.1947 |
| Intercept + Temperature + Precipitation + Admixture:Precipitation | 4 | 0.2528 | 0.1939 |
| Intercept + Temperature + Precipitation | 3 | 0.2327 | 0.1933 |
| Intercept + Temperature + Temperature:Precipitation | 3 | 0.2312 | 0.1918 |
| Intercept + Temperature + Precipitation + Admixture:Temperature:Precipitation | 4 | 0.2508 | 0.1916 |
| Intercept + Temperature | 2 | 0.2111 | 0.1914 |
| Intercept + Temperature + Admixture:Precipitation + Temperature:Precipitation | 4 | 0.2500 | 0.1908 |
| Intercept + Temperature + Admixture:Temperature:Precipitation | 3 | 0.2302 | 0.1907 |
| Intercept + Temperature + Admixture:Precipitation | 3 | 0.2260 | 0.1863 |
| Intercept + Admixture:Temperature + Admixture:Precipitation + Temperature:Precipitation + Admixture:Temperature:Precipitation | 5 | 0.2341 | 0.1513 |
| Intercept + Admixture:Temperature + Admixture:Precipitation + Admixture:Temperature:Precipitation | 4 | 0.2098 | 0.1475 |
| Intercept + Precipitation + Admixture:Temperature + Admixture:Precipitation + Admixture:Temperature:Precipitation | 5 | 0.2280 | 0.1446 |
| Intercept + Precipitation + Admixture:Temperature + Admixture:Precipitation + Temperature:Precipitation + Admixture:Temperature:Precipitation | 6 | 0.2486 | 0.1442 |
| Intercept + Admixture:Precipitation + Temperature:Precipitation + Admixture:Temperature:Precipitation | 4 | 0.1951 | 0.1315 |
| Intercept + Precipitation + Admixture:Precipitation + Temperature:Precipitation + Admixture:Temperature:Precipitation | 5 | 0.2104 | 0.1250 |
| Intercept + Precipitation + Admixture:Temperature + Admixture:Precipitation | 4 | 0.1881 | 0.1240 |
| Intercept + Precipitation + Admixture:Precipitation + Admixture:Temperature:Precipitation | 4 | 0.1869 | 0.1227 |
| Intercept + Admixture:Precipitation + Admixture:Temperature:Precipitation | 3 | 0.1633 | 0.1204 |
| Intercept + Precipitation + Admixture:Temperature + Admixture:Precipitation + Temperature:Precipitation | 5 | 0.2058 | 0.1199 |
| Intercept + Precipitation + Admixture:Temperature + Temperature:Precipitation + Admixture:Temperature:Precipitation | 5 | 0.2029 | 0.1167 |
| Intercept + Precipitation + Admixture:Temperature + Temperature:Precipitation | 4 | 0.1792 | 0.1144 |
| Intercept + Admixture:Temperature + Temperature:Precipitation + Admixture:Temperature:Precipitation | 4 | 0.1783 | 0.1134 |
| Intercept + Precipitation + Admixture:Temperature | 3 | 0.1498 | 0.1062 |
| Intercept + Precipitation + Admixture:Temperature + Admixture:Temperature:Precipitation | 4 | 0.1706 | 0.1051 |
| Intercept + Admixture:Temperature + Admixture:Temperature:Precipitation | 3 | 0.1487 | 0.1050 |
| Intercept + Precipitation + Admixture:Precipitation + Temperature:Precipitation | 4 | 0.1673 | 0.1016 |
| Intercept + Precipitation + Admixture:Precipitation | 3 | 0.1447 | 0.1008 |
| Intercept + Precipitation + Temperature:Precipitation + Admixture:Temperature:Precipitation | 4 | 0.1641 | 0.0981 |
| Intercept + Precipitation + Temperature:Precipitation | 3 | 0.1390 | 0.0948 |
| Intercept + Temperature:Precipitation + Admixture:Temperature:Precipitation | 3 | 0.1375 | 0.0932 |
| Intercept + Precipitation | 2 | 0.1074 | 0.0851 |
| Intercept + Precipitation + Admixture:Temperature:Precipitation | 3 | 0.1286 | 0.0839 |
| Intercept + Admixture:Temperature:Precipitation | 2 | 0.1034 | 0.0810 |
| Intercept + Admixture:Temperature | 2 | 0.0508 | 0.0270 |
| Intercept + Admixture:Temperature + Temperature:Precipitation | 3 | 0.0720 | 0.0245 |
| Intercept + Admixture:Temperature + Admixture:Precipitation | 3 | 0.0705 | 0.0228 |
| Intercept + Admixture:Temperature + Admixture:Precipitation + Temperature:Precipitation | 4 | 0.0892 | 0.0173 |
| Intercept + Temperature:Precipitation | 2 | 0.0317 | 0.0075 |
| Intercept + Admixture:Precipitation + Temperature:Precipitation | 3 | 0.0499 | 0.0012 |
| Intercept + Admixture:Precipitation | 2 | 0.0248 | 0.0004 |
| Intercept | 1 | 0.0000 | 0.0000 |

Table S3: Partial $R^2$ for predictors included in the best-fit genotype-environment-association model. RSS is residual sums of squares for the best-fit ($RSS_{best}$) and reduced ($RSS_{reduced}$) multivariate-normal linear regression models. For each predictor, the reduced model is the best-fit model with the predictor omitted. Predictors are sorted by decreasing partial $R^2$.

| Predictor | $RSS_{reduced}$ | $RSS_{best}$ | Partial $R^2$ |
|---|---|---|---|
| Admixture | 30339.0 | 23405.2 | 0.2285 |
| Temperature | 25765.5 | 23405.2 | 0.0916 |
| Admixture:Temperature | 25252.0 | 23405.2 | 0.0731 |
| Admixture:Precipitation | 24421.3 | 23405.2 | 0.0416 |
| Admixture:Temperature:Precipitation | 24283.0 | 23405.2 | 0.0362 |
| Precipitation | 24128.9 | 23405.2 | 0.0300 |

Table S4: Gene-enrichment tests for each predictor in the best-fit genotype-environment-association (GEA) model. Across all SNPs in the dataset, 84.7% were annotated as gene-associated. For each predictor in the best-fit GEA model, the proportion of statistically-significant SNPs which were gene-associated (false-discovery-rate $\alpha = 0.05$; Significant) was compared to the overall proportion of SNPs in the study which were gene-associated (Ratio). The $p$-values from non-parametric resampling analyses test whether the proportion of statistically-significant SNPs which were gene-associated was higher than expected from random sampling. *Statistical significance at $\alpha = 0.05$.

| | Gene-associated | | | |
|---|---|---|---|---|
| Predictor | Significant | Overall | Ratio | $p$-value |
| Admixture | 0.859 | 0.847 | 1.014 | $< 0.001^*$ |
| Temperature | 0.857 | 0.847 | 1.012 | $< 0.001^*$ |
| Precipitation | 0.846 | 0.847 | 0.999 | 0.587 |
| Admixture:Temperature | 0.862 | 0.847 | 1.017 | $< 0.001^*$ |
| Admixture:Precipitation | 0.861 | 0.847 | 1.016 | $0.017^*$ |
| Admixture:Temperature:Precipitation | 0.849 | 0.847 | 1.003 | 0.365 |

Table S5: Statistically-significant gene-ontology-enrichment tests for each predictor in the best-fit genotype-environment-association (GEA) model. For each combination of predictor and gene ontology number, the proportion of statistically-significant SNPs which were annotated with the gene ontology number (false-discovery-rate [FDR] $\alpha = 0.05$; Significant) were compared to the overall proportion of SNPs in the study which were annotated with the gene ontology number (Ratio). The $p$-values from non-parametric resampling analyses test whether the proportion of statistically-significant SNPs which were annotated with the gene ontology number was higher than expected from random sampling. *FDR-adjusted $p$-values. All displayed tests were statistically-significant at $\alpha = 0.05$.

| | Gene ontology | | Gene ontology annotation | | | |
|---|---|---|---|---|---|---|
| Predictor | Number | Description | Significant | Overall | Ratio | $p$-value* |
| Admixture | #0005509 | Molecular function: Calcium ion binding | 0.01140 | 0.00937 | 1.216 | < 0.001 |
| Admixture | #0005515 | Molecular function: Protein binding | 0.07186 | 0.06600 | 1.089 | < 0.001 |
| Admixture | #0030008 | Cellular component: TRAPP complex | 0.00088 | 0.00042 | 2.079 | < 0.001 |
| Temperature | #0004174 | Molecular function: Electron-transferring-flavoprotein dehydrogenase activity | 0.00070 | 0.00014 | 5.016 | 0.034 |
| Temperature | #0004739 | Molecular function: Pyruvate dehydrogenase (acetyl-transferring) activity | 0.00055 | 0.00011 | 5.201 | < 0.001 |
| Temperature | #0006086 | Biological process: Acetyl-CoA biosynthetic process from pyruvate | 0.00055 | 0.00011 | 5.201 | 0.034 |
| Temperature | #0022900 | Biological process: Electron transport chain | 0.00070 | 0.00017 | 4.180 | < 0.001 |
| Temperature | #0090630 | Biological process: Activation of GTPase activity | 0.00094 | 0.00030 | 3.110 | 0.049 |
| Temperature | #0106035 | Biological process: Protein maturation by [4Fe-4S] cluster transfer | 0.00055 | 0.00010 | 5.573 | 0.049 |
| Temperature | #1990316 | Cellular component: Atg1/ULK1 kinase complex | 0.00055 | 0.00009 | 6.001 | < 0.001 |
| Precipitation | #0004563 | Molecular function: $\beta$-N-acetylhexosaminidase activity | 0.00269 | 0.00034 | 7.813 | < 0.001 |
| Admixture:Temperature | #1990316 | Cellular component: Atg1/ULK1 kinase complex | 0.00071 | 0.00009 | 7.718 | < 0.001 |
| Admixture:Temperature:Precipitation | #0000289 | Biological process: Nuclear-transcribed mRNA poly(A) tail shortening | 0.00124 | 0.00006 | 19.606 | < 0.001 |
| Admixture:Temperature:Precipitation | #0004563 | Molecular function: $\beta$-N-acetylhexosaminidase activity | 0.00217 | 0.00034 | 6.302 | < 0.001 |
| Admixture:Temperature:Precipitation | #0031251 | Cellular component: PAN complex | 0.00124 | 0.00006 | 19.606 | < 0.001 |

### Supplemental Figures

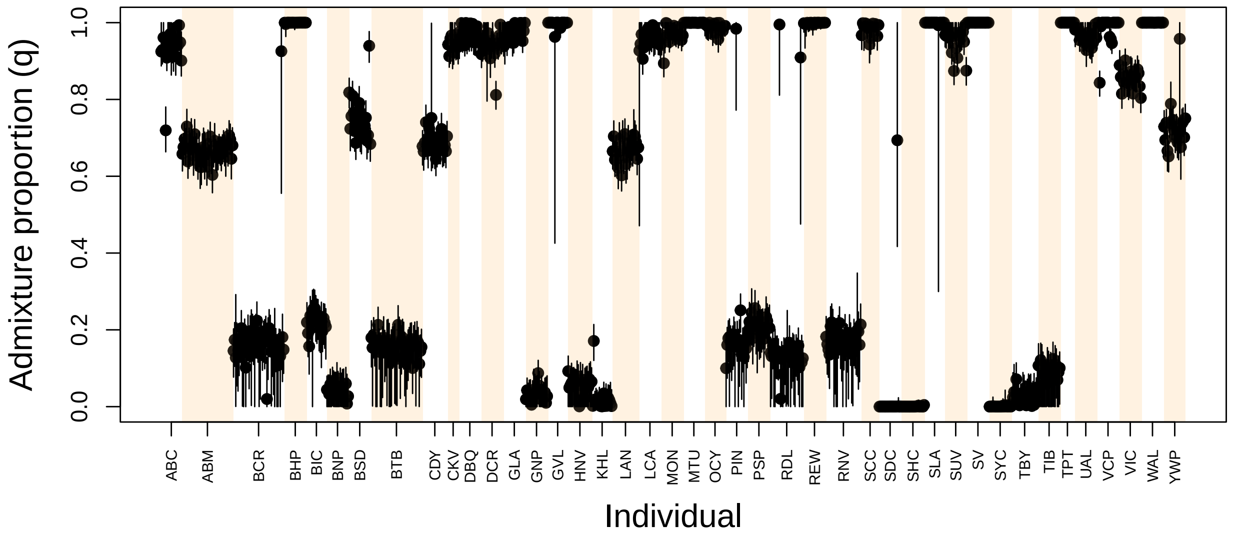


Figure S1: Admixture proportions from the 922 *Lycaeides* butterflies. Points and vertical line segments denote Bayesian estimates of admixture proportions ($q$) and 95% equal-tail probability intervals, respectively. The 922 individual butterflies are sorted by population, with populations organized alphabetically (colored shading delineates populations on the plot). See Table 1 for detailed population information.

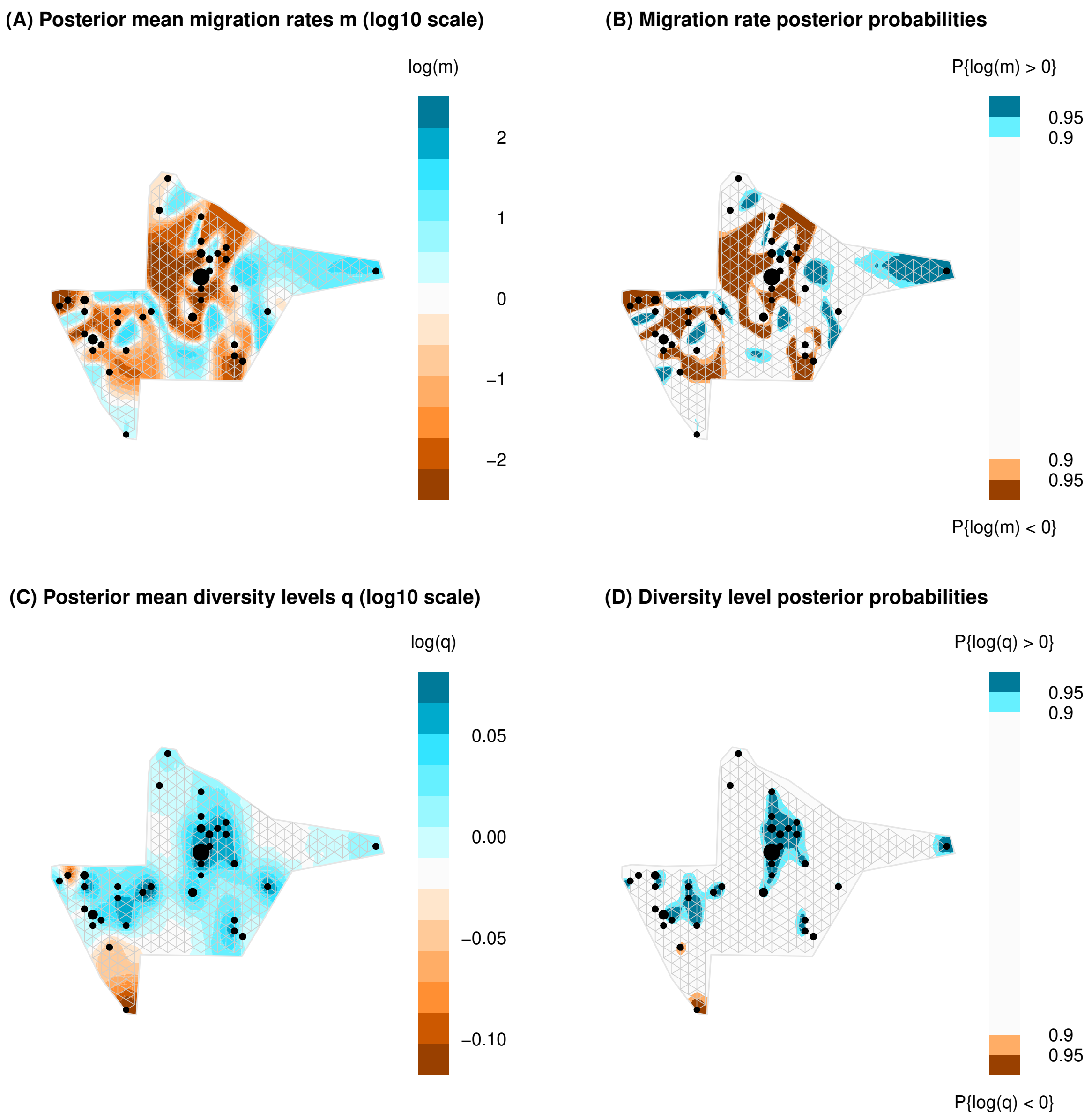


Figure S2: Relative effective migration and effective diversity surfaces. Examples are shown for one of three Markov chain Monte Carlo runs with 400 deme. Surfaces are colored to denote relative migration rates (A), relative migration rate posterior probabilities (B), relative diversity levels (C), and relative diversity level posterior probabilities (D). The triangular grid that defines deme locations is given in each panel. Points denote sampled butterflies with circle sizes proportional to sample sizes. In some cases, butterflies from nearby locations were treated as a single deme based on the grid.

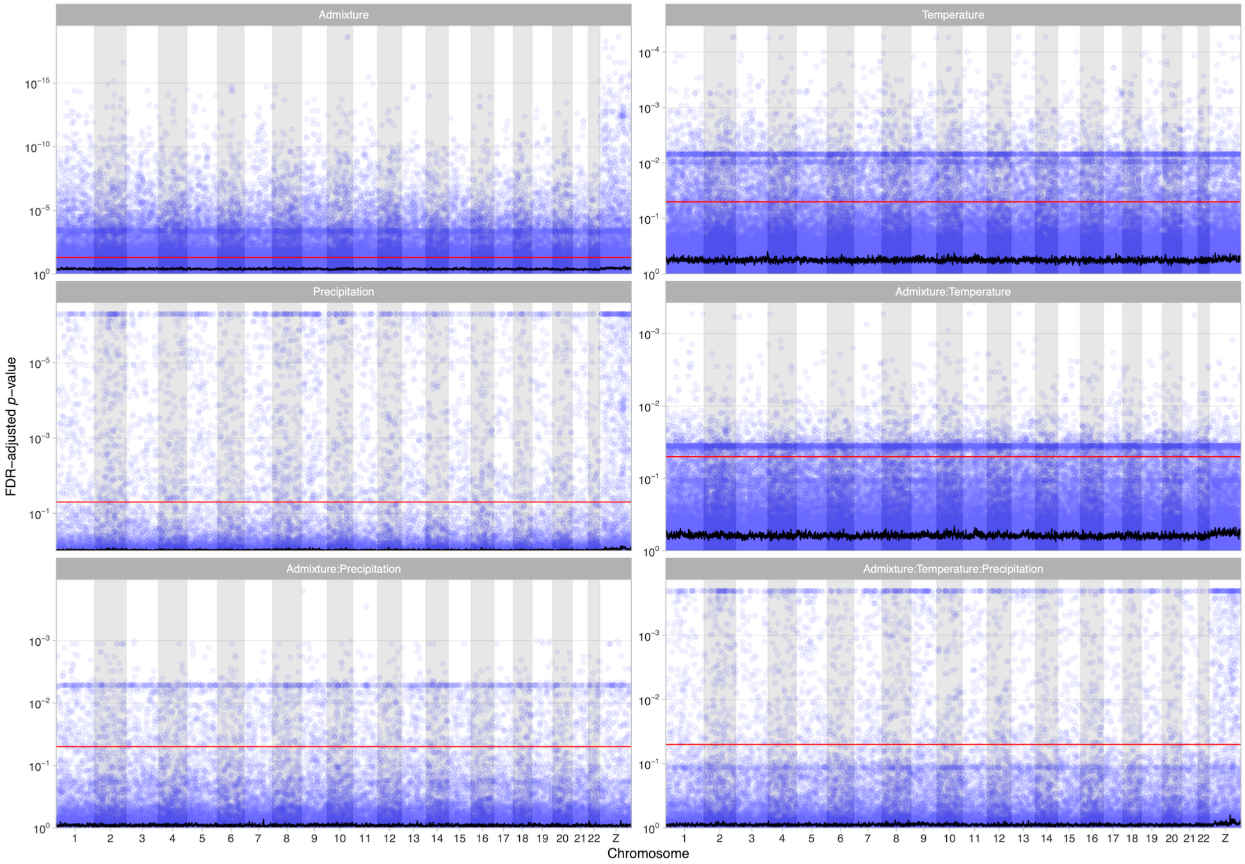


Figure S3: False-discovery-rate (FDR) adjusted $p$-values for each SNP in the best-fit genotype-environment-association model by position along each chromosome. The horizontal red line represents an $\alpha$ level of 0.05. The black line represents a moving average window of 100 SNPs. Horizontal banding of points is a byproduct of the FDR adjustments. Note the $\log_{10}$ scale on the y-axis.
